# Control of Synchrony by Inhibitory Motifs in Spiking Neural Networks

**DOI:** 10.1101/2025.11.07.687261

**Authors:** Archishman Biswas, Arvind Kumar

**Affiliations:** School of Electrical Engineering and Computer Science KTH Royal Institute of Technology, Stockholm, Sweden; SciLifeLab, Solna, Sweden

**Keywords:** Non-random networks, EI networks, Synchrony, Triplet Motifs

## Abstract

Connectivity in biological neuronal networks is significantly different from the random network model (Erdős-Rényi). Specifically, di-synaptic motifs like reciprocal, convergent, divergent, and chain are found to be either over-represented or under-represented in certain brain regions. Over-representation of such motifs among excitatory neurons is known to induce synchrony. However, cortical activity is typically asynchronous. Thus, it remains unclear how synchrony induced by excitatory motifs may be reduced to physiological levels. To address this question, we systematically vary the prevalence of these four motifs in an Excitatory-Inhibitory (EI) network. We found that over-representation of chain and convergent motifs in the excitatory population led to increased firing rates and greater synchrony. However, this excess synchrony was quenched when we introduced the same type of motifs among inhibitory neurons. Because of the overabundance of motifs, some inhibitory neurons received fewer recurrent inhibitory inputs. Such weakly coupled neurons were primarily driven by uncorrelated external inputs, and therefore, these neurons exerted stronger inhibition on excitatory neurons and reduced both synchrony and firing rates. Thus, we also provide a new mechanism by which synchrony can be controlled in excitatory-inhibitory networks. We predict that the same kind of di-synaptic motifs should exist in both excitatory and inhibitory neurons.

**Significance Statement:** Asynchronous cortical activity is inconsistent with the overabundance of di-synaptic motifs (e.g. convergent, chain, etc.). In this work, we resolve this inconsistency. We show that inducing di-synaptic motifs in the inhibitory population can quench the synchrony induced by such motifs among excitatory neurons. We show that di-synaptic motifs give rise to high variance in neurons’ degree distribution. We found that inhibitory neurons with fewer in-degree are largely driven by external inputs; therefore, they spike in an uncorrelated manner at high firing rate. Such neurons decorrelate the network activity. Thus, in addition to reconciling the mismatch between network structure and network activity, we reveal a new mechanism to control synchrony in biological neuronal networks.

## Introduction

Understanding the relationship between network structure and network activity dynamics is a fundamental question in neuroscience. In general, neuronal network activity depends upon input signals, neuron dynamics, synapse dynamics, and connectivity structure. The connectivity structure of cortical networks depends on the spatial scales. At macroscopic scales, the inter-region connectivity is highly non-random and tends to exhibit small-world or scale-free properties [1, 2, 3, 4]. At the scale of 1 *mm*, the connection probability monotonically decays with distance between neurons [5, 6]. Zooming in further at the scale of *<* 100 *µm* experimental data shows that 3-neuron motifs (triplet motifs) are either overrepresented or underrepresented in cortical micro-circuits [7, 8, 9, 10]. Recent data from the FlyWire project also suggest that networks in the fly brain have non-random connectivity in different regions of the Drosophila brain, even at very small spatial scales [11].

Effects of connectivity features that are observed *<* 1 mm scale is rather well understood. For instance, local connectivity [12, 13, 14, 15, 16], global matrix features e.g. low-rankness [17, 18, 19] and degree distributions [20, 21] can predict the activity state of neuronal networks. By contrast, much less is understood about how motifs of 3-10 neurons shape the network structure and thereby affect the network activity dynamics. Previous work suggests that a random network with an abundance of di-synaptic divergent, convergent or chain type motifs tend to synchronize and oscillate more than a corresponding Erdő s–Rényi (ER) type network [22, 23, 16]. Also, network features such as degree distributions (which are related to di-synaptic motifs) can affect synchrony and oscillations in EI networks [20]. However, such excessive synchrony is inconsistent with the experimental data on network activity *in vivo*, which is typically characterized by an asynchronous and irregular state [24]. Therefore, the question arises: how is the synchrony induced by these di-synaptic motifs among excitatory neurons quenched?

To address this question, we systematically varied the proportion of reciprocal, convergent, divergent, and chain motifs in either E or I or both E and I populations. At the structural level, an abundance of convergent and divergent motifs broadened the in-degree and out-degree distributions, respectively, relative to those of Erdő s–Rényi (ER) random networks. Chain motifs, in addition, introduced correlation between in- and out-degree distributions. Higher chain motifs than the ER baseline led to positive correlations between the in- and out-degree distributions, whereas lower chain motifs than the ER baseline led to negative correlations. In our simulations, only the convergent and chain motifs in the E population substantially affected the dynamics, leading to higher firing rates and stronger synchrony. By contrast, reciprocal and divergent motifs had negligible effects. On the other hand abundance of convergent and chain motifs in the inhibitory population reduced the firing rate and synchrony. When excess motifs are present in both populations, inhibitory motifs (chain and convergent type) effectively cancel the synchrony induced by excitatory motifs. As a consequence of di-synaptic motifs, the in-degree distribution variance increases, and some inhibitory neurons end up receiving rather few recurrent inhibitory inputs. It is these weakly recurrent inhibitory neurons that cancel the synchrony induced by excitatory motifs. While inhibitory motifs can cancel the synchrony, overall, the interaction between of excitatory and inhibitory motifs significantly expands the dynamical range of network activity.

We predict that in the cortical EI networks, there shall be a sufficient number of inhibitory convergent and chain motifs to counter the synchrony-promoting effects of the same motifs in the excitatory population. This helps in maintaining an irregular-asynchronous state that is observed in *in vivo*.

## Methods

### Neuron Model

The neurons were modeled as leaky integrate-and-fire (LIF) neurons. The equation defining the sub-threshold dynamics of a LIF neuron is given as follows:

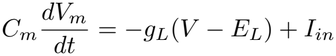

where *C*_*m*_ is membrane capacitance, *E*_*L*_ is the leak reversal potential and *g*_*L*_ is the leak conductance. When the membrane potential reached a threshold, i.e. *V*_*m*_ *≥ V*_*th*_, the neuron spiked, and the membrane potential was reset to a value *V*_*m*_ = *V*_*r*_ (reset potential) for a refractory period of *t*_*r*_.

To test whether the effects of di-synaptic motifs depended on the neuronal model, we repeated selected simulations using exponential integrate-and-fire (EIF) neurons. Their subthreshold membrane-potential dynamics were described by

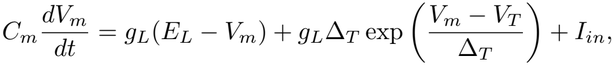

where *V*_*T*_ is the effective threshold of the exponential current and Δ_*T*_ determines the sharpness of spike initiation. When the membrane potential crossed the spike-detection threshold, *V*_*m*_ *≥ V*_spike_, a spike was recorded and the membrane potential was reset to value *V*_*m*_ = *V*_*th*_ for a refractory period of *t*_*r*_. The EIF model therefore differed from the LIF model through the additional exponential term describing nonlinear spike initiation.

The input current *I*_*in*_ into the LIF neuron consisted of three parts:

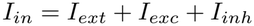

where *I*_*exc*_ and *I*_*inh*_ are the excitatory currents and inhibitory currents due to synaptic input from other excitatory and inhibitory neurons in the network, respectively. The external input was modeled as Poisson type spike trains; therefore, *I*_*ext*_ had the same form as that of excitatory currents *I*_*exc*_.

### Synapse Model

Synapses were modeled as conductance-transients. Each incoming spikes created a conductance transient *g*_*syn*_(*t*):

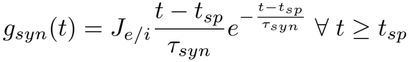

where *syn ∈* [*exc, inh*], *t*_*sp*_ is time of incoming spike, *τ*_*syn*_ is the synaptic time constant and, *J*_*syn*_ is the peak conductance. The total excitatory or inhibitory currents coming into a neuron will be the sum of all the conductance transient effects from the pre-synaptic neurons:

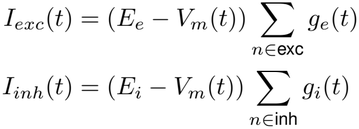

In the equations for excitatory and inhibitory currents, the summation is across all excitatory and inhibitory synapses that project upon a neuron, respectively. The *E*_*e*_ and *E*_*i*_ are the reversal potentials for the excitatory and inhibitory synapses. For all our simulation studies, we chose the *J*_*e*_ such that the excitatory post-synaptic potential (ePSP) turns out to be around 0.1 *mV* when the post-neuron was at a resting potential of *−*70 *mV* . And the *J*_*i*_ such that the inhibitory post-synaptic potential (iPSP) turns out to be around *−*0.75 *mV* when the post-neuron was at a resting potential of *−*55 *mV* . Hence, the ratio (*γ*) of inhibitory to excitatory PSPs was set around 7.5.

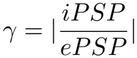

### Input to the Network

We simulated the networks by injecting independent Poisson spike trains into each neuron. The parameters of the input are the rate of the Poisson process and the strength of the synaptic connectivity it makes at the target neurons. Each of the neurons was injected with a different realization of the Poisson spike train. The input rate of the spikes was set to 1000 *Hz* and the synaptic parameter *J*_*e*_ was chosen such that the ePSP from external synapses turns out to be around 1.0 *mV* . The associated parameters with the EI network are also shown in the figure 1C and are listed in table 1 (see table 2 for parameters used to simulate networks with EIF neuron model).

**Table 1:**
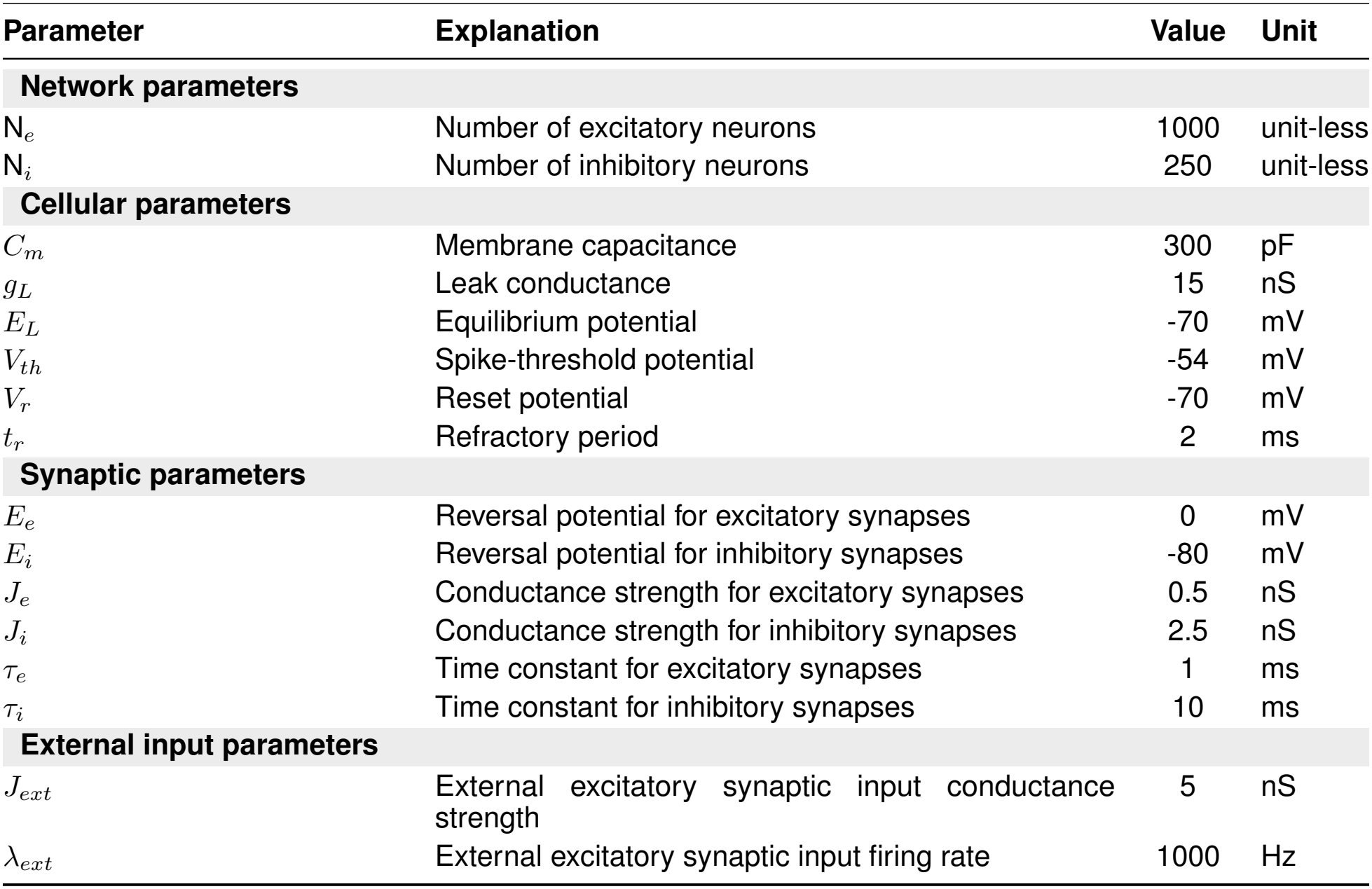
Parameters used in the LIF neuronal network simulations.

| Parameter | Explanation | Value | Unit |
|---|---|---|---|
| **Network parameters** |  |  |  |
| N<sub>*e*</sub> | Number of excitatory neurons | 1000 | unit-less |
| N<sub>*i*</sub> | Number of inhibitory neurons | 250 | unit-less |
| **Cellular parameters** |  |  |  |
| *C<sub>m</sub>* | Membrane capacitance | 300 | pF |
| *g<sub>L</sub>* | Leak conductance | 15 | nS |
| *E<sub>L</sub>* | Equilibrium potential | -70 | mV |
| *V<sub>th</sub>* | Spike-threshold potential | -54 | mV |
| *V<sub>r</sub>* | Reset potential | -70 | mV |
| *t<sub>r</sub>* | Refractory period | 2 | ms |
| **Synaptic parameters** |  |  |  |
| *E<sub>e</sub>* | Reversal potential for excitatory synapses | 0 | mV |
| *E<sub>i</sub>* | Reversal potential for inhibitory synapses | -80 | mV |
| *J<sub>e</sub>* | Conductance strength for excitatory synapses | 0.5 | nS |
| *J<sub>i</sub>* | Conductance strength for inhibitory synapses | 2.5 | nS |
| *τ<sub>e</sub>* | Time constant for excitatory synapses | 1 | ms |
| *τ<sub>i</sub>* | Time constant for inhibitory synapses | 10 | ms |
| **External input parameters** |  |  |  |
| *J<sub>ext</sub>* | External excitatory synaptic input conductance strength | 5 | nS |
| *λ<sub>ext</sub>* | External excitatory synaptic input firing rate | 1000 | Hz |

**Table 2:** Parameters used in the EIF neuronal network simulations.

| Parameter | Explanation | Value | Unit |
|---|---|---|---|
| **Network parameters** |  |  |  |
| *N*<sub>*e*</sub> | Number of excitatory neurons | 1000 | unit-less |
| *N*<sub>*i*</sub> | Number of inhibitory neurons | 250 | unit-less |
| **Cellular parameters** |  |  |  |
| *C*<sub>*m*</sub> | Membrane capacitance | 300 | pF |
| *g*<sub>*L*</sub> | Leak conductance | 15 | nS |
| *E*<sub>*L*</sub> | Leak reversal potential | -70 | mV |
| *V*<sub>*T*</sub> | Effective threshold of the exponential current | -60 | mV |
| Δ<sub>*T*</sub> | Sharpness of spike initiation | 3 | mV |
| *V*<sub>*r*</sub> | Reset potential | -70 | mV |
| *V*<sub>spike</sub> | Spike-detection threshold | -40 | mV |
| *t*<sub>*r*</sub> | Refractory period | 2 | ms |
| **Synaptic parameters** |  |  |  |
| *E*<sub>*e*</sub> | Reversal potential for excitatory synapses | 0 | mV |
| *E*<sub>*i*</sub> | Reversal potential for inhibitory synapses | -80 | mV |
| *J*<sub>*e*</sub> | Conductance strength for excitatory synapses | 0.5 | nS |
| *J*<sub>*i*</sub> | Conductance strength for inhibitory synapses | 2.5 | nS |
| *τ*<sub>*e*</sub> | Time constant for excitatory synapses | 1 | ms |
| *τ*<sub>*i*</sub> | Time constant for inhibitory synapses | 10 | ms |
| **External input parameters** |  |  |  |
| *J*<sub>ext</sub> | External excitatory synaptic conductance strength | 5 | nS |
| *λ*<sub>ext</sub> | External excitatory input firing rate | 750 | Hz |

**Figure 1:**
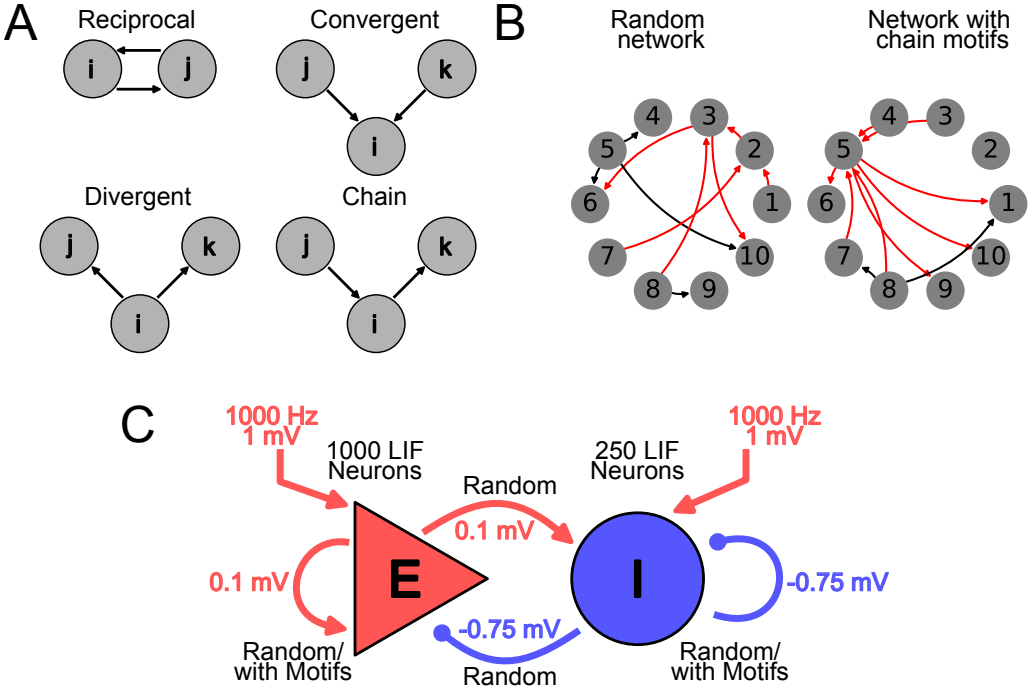
Di-synaptic motifs and schematic of the excitatory-inhibitory (EI) network model. (**A**) Schematic of the four triplet motifs, namely reciprocal, convergent, divergent, and chain, whose effects on neuronal network dynamics were investigated. (**B**) Two illustrative example networks with ten neurons. Left: Network with random connectivity. Right: Network with more chain motifs than the random network baseline. The edges in red contribute to the total count of chain motifs in the network. (**C**) Schematic of the EI network showing the associated number of neurons, synaptic weights, inputs, and type of connectivity.

### Heterogeneous Recruitment of the Inhibitory Population

In the experiments shown in 4A, to introduce heterogeneity without changing the inhibitory connectivity, we randomly selected 75% of the inhibitory neurons in networks with E-chain/I-random connectivity. The selected neurons received an additional constant current. Within each simulation condition, the same current amplitude was applied to every selected neuron. Across conditions, the current amplitude was varied from 0 pA to 120 pA in increments of 20 pA. Ten independent trials were performed for each current amplitude.

### Random Network Model

The network model as shown in 1C consists of 1000 excitatory(E) neurons and 250 inhibitory(I) neurons (EI network). The connectivity probability from a source (*s*) population to a target (*t*) population is given by *p*_*ts*_. For the random network case, all the connections between the excitatory and the inhibitory population had a probability of *p*_*ee*_ = *p*_*ie*_ = *p*_*ei*_ = *p*_*ii*_ = 0.1. This network is a random network or an Erdő s–Rényi(ER) network as the probability of any pair of neurons being connected is the same. Two within-population connectivity matrices in this network connect the *E → E* and *I → I*. These two connectivity matrices that connect the E or I population to itself are denoted by ***W***_***EE***_ and ***W***_***II***_, respectively. Later, when we describe networks with non-ER connectivity, we only made modifications to these square matrices ***W***_***EE***_ and ***W***_***II***_ . And, we do not change the ***W***_***IE***_ and ***W***_***EI***_ that connect the *E → I* and *I → E*, respectively.

### Parameters Used in Network Simulations

Here we give a list of various parameters that were used in the network simulations, along with the explanation, values, and units (table 1). For the figure S7 A, the *λ*_*ext*_ was varied over different ranges in the different representative networks to get the desired population firing rate.

### Generation of Connectivity Matrix with Fixed Motifs Statistics

The experimental observations show the prevalence of different triplet motifs (3-node sub-networks) in brain microcircuits. A simpler subset of these motifs consists of only two synaptic edges, or second order motifs. These motifs are reciprocal, convergent, divergent, and chain [22] as shown in figure 1A. If we think about the edges of the neuronal network, then these four quantities are related to the correlation between the edges. For a network defined by a fixed fraction of these four motifs, we need to add four more parameters (one for each of the motifs) in addition to the sparsity parameter *p*, which is a parameter for the random networks already. These four parameters are *α*_*recip*_, *α*_*conv*_, *α*_*div*_, *α*_*chain*_. As shown in the work [22], these are related to the joint probabilities of different pairs of edges in a connectivity matrix ***W*** as follows:

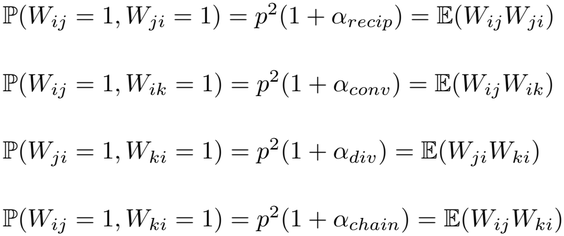

For a chosen set of *α* values, four equations set joint constraints on different entries of the adjacency matrix ***W*** _*ij*_. And, we also have the constraint set by the sparsity parameter *p* of the network as follows:

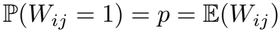

The SONET algorithm is used to generate connectivity matrices ***W*** based on the constraints set by the five parameters *p, α*_*recip*_, *α*_*conv*_, *α*_*div*_, *α*_*chain*_. These parameters cannot be chosen completely independently of each other. Only a small set of parameters allows the creation of a valid connectivity network. For a chosen average probability of connection *p*, the parameter *α*_*recip*_ can range in *−*1 to 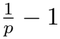. The *α*_*conv*_ and *α*_*div*_ must be non-negative as they represent variances in the in and out degree distributions, respectively. Also, their maximum value can be 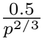. And, finally for a chosen *α*_*conv*_ and *α*_*div*_, the *α*_*chain*_ lies in the range 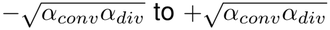. Hence, the total set of usable parameters is quite restrictive. But, for our purpose, we can create connectivity matrices that have enough non-randomness (i.e. deviations w.r.t. ER networks) to affect the dynamics even with this restrictive set of parameters. Examples of the matrices generated by the algorithm are shown in the figure 2(A). By visualizing the matrices generated by SONET, we can see that even with the restrictive set of parameters, the generated matrices show the non-random patterns prominently. The average probability of connection (sparsity *p*) was set to 0.1 for each case.

**Figure 2:**
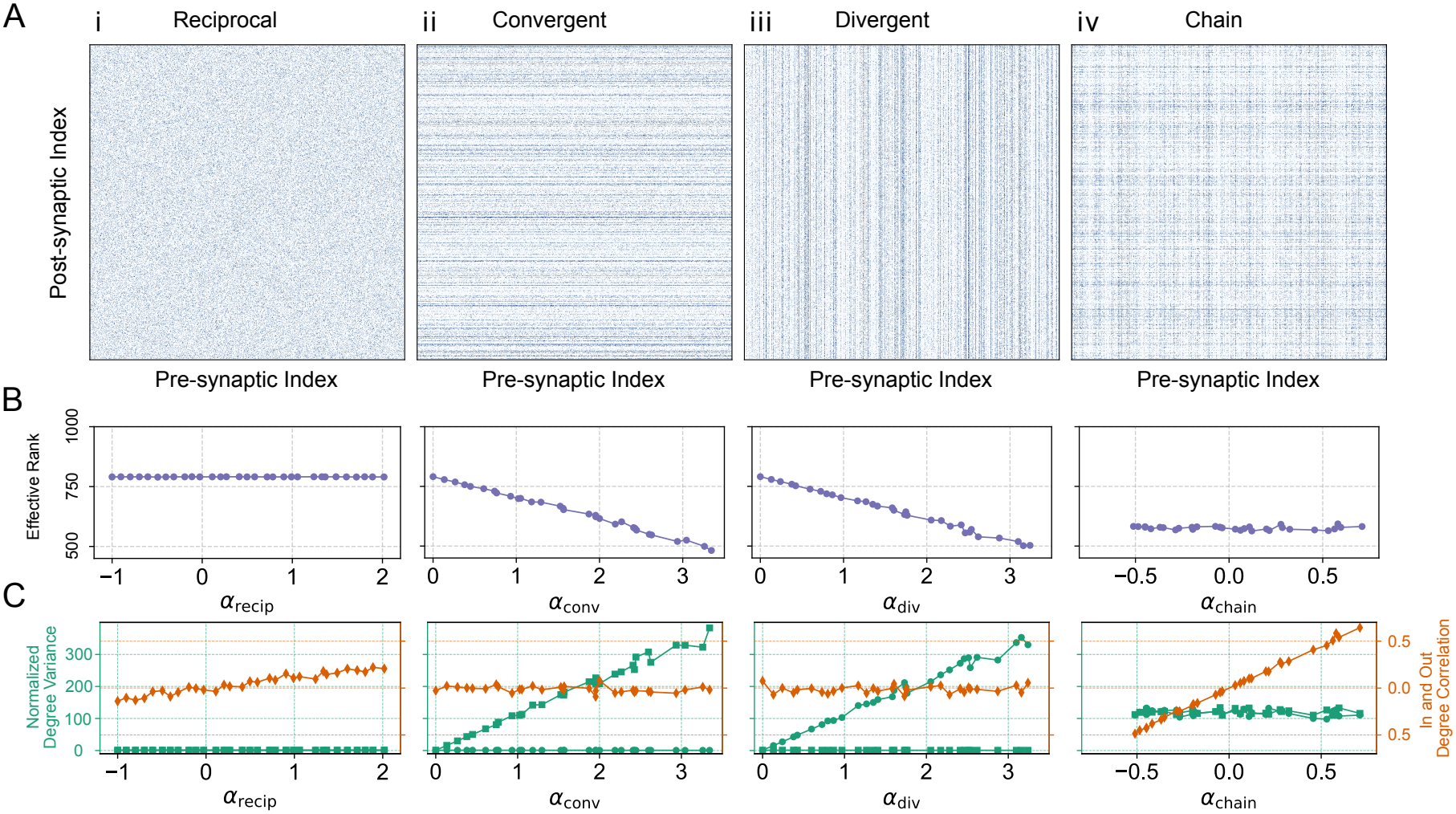
Di-synaptic motifs affect higher order properties of the connectivity matrix. (**A**) Examples of connectivity matrices of different *α* values were generated using the SONETs algorithm. The target (post-synaptic) neurons are shown in the vertical axis, and the source (pre-synaptic) neurons are shown in the horizontal axis. The average probability of connection (sparsity parameter *p*) was set to 0.1 for all four networks. (**B**) Effective ranks of the connectivity matrices with different kinds of motifs. For the first three plots from the left, only the *α*_*recip*_, *α*_*conv*_, *α*_*div*_ were varied, respectively, while other *α* values were set to zero. For the last plot, the *α*_*recip*_ was set to 0, *α*_*conv*_ and *α*_*div*_ were set to 1.0, and then the *α*_*chain*_ was varied. Sparsity was set to *p* = 0.1 for all the networks. (**C**) The *α* values were varied in the same fashion as those of the plots in panel **B**, and also sparsity was set to *p* = 0.1. The left axis of the plots shows the variance of the in-degree (in square markers) and out-degree (in circular markers) distribution. Variances were normalized with respect to the variances in an ER network with the same average connection probability (*p* = 0.1). The right axis displays the correlation between the in-degree and out-degree distributions.

To clarify how the addition of motifs changes the network architecture, we illustrate the case of chain motifs. Increasing the prevalence of these motifs does not change the total number of synapses but rearranges them so that the neurons receiving more input synapses also send more output synapses (Figure 1B). Hence, we can change the network structure without changing the density or average probability of connections.

### Varying *α* Parameters

To study the effect of the prevalence of different motifs, we had set the parameters *α*_*recip*_, *α*_*conv*_, *α*_*div*_, *α*_*chain*_ to different values for different simulations. This section gives the details on how the parameters were varied. The *α* values are also summarized in the table 3.

**Table 3:**
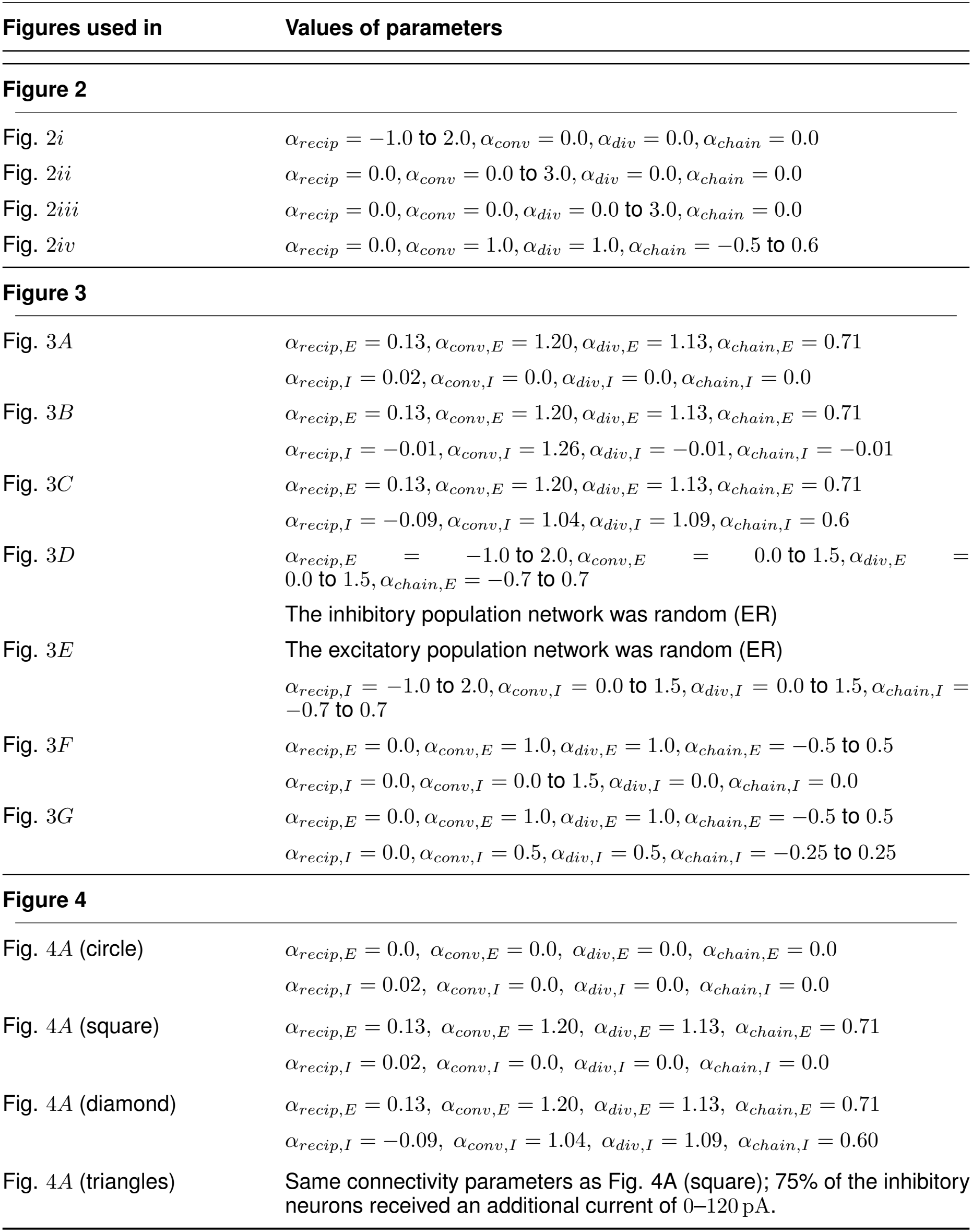
Values of the *α* parameters used in different simulations.

| Figures used in | Values of parameters |
|---|---|
| **Figure 2** |  |
| Fig. 2*i* | α<sub>recip</sub> = −1.0 to 2.0, α<sub>conv</sub> = 0.0, α<sub>div</sub> = 0.0, α<sub>chain</sub> = 0.0 |
| Fig. 2*ii* | α<sub>recip</sub> = 0.0, α<sub>conv</sub> = 0.0 to 3.0, α<sub>div</sub> = 0.0, α<sub>chain</sub> = 0.0 |
| Fig. 2*iii* | α<sub>recip</sub> = 0.0, α<sub>conv</sub> = 0.0, α<sub>div</sub> = 0.0 to 3.0, α<sub>chain</sub> = 0.0 |
| Fig. 2*iv* | α<sub>recip</sub> = 0.0, α<sub>conv</sub> = 1.0, α<sub>div</sub> = 1.0, α<sub>chain</sub> = −0.5 to 0.6 |
| **Figure 3** |  |
| Fig. 3*A* | α<sub>recip,E</sub> = 0.13, α<sub>conv,E</sub> = 1.20, α<sub>div,E</sub> = 1.13, α<sub>chain,E</sub> = 0.71 |
|  | α<sub>recip,I</sub> = 0.02, α<sub>conv,I</sub> = 0.0, α<sub>div,I</sub> = 0.0, α<sub>chain,I</sub> = 0.0 |
| Fig. 3*B* | α<sub>recip,E</sub> = 0.13, α<sub>conv,E</sub> = 1.20, α<sub>div,E</sub> = 1.13, α<sub>chain,E</sub> = 0.71 |
|  | α<sub>recip,I</sub> = −0.01, α<sub>conv,I</sub> = 1.26, α<sub>div,I</sub> = −0.01, α<sub>chain,I</sub> = −0.01 |
| Fig. 3*C* | α<sub>recip,E</sub> = 0.13, α<sub>conv,E</sub> = 1.20, α<sub>div,E</sub> = 1.13, α<sub>chain,E</sub> = 0.71 |
|  | α<sub>recip,I</sub> = −0.09, α<sub>conv,I</sub> = 1.04, α<sub>div,I</sub> = 1.09, α<sub>chain,I</sub> = 0.6 |
| Fig. 3*D* | α<sub>recip,E</sub> = −1.0 to 2.0, α<sub>conv,E</sub> = 0.0 to 1.5, α<sub>div,E</sub> = 0.0 to 1.5, α<sub>chain,E</sub> = −0.7 to 0.7 |
|  | The inhibitory population network was random (ER) |
| Fig. 3*E* | The excitatory population network was random (ER) |
|  | α<sub>recip,I</sub> = −1.0 to 2.0, α<sub>conv,I</sub> = 0.0 to 1.5, α<sub>div,I</sub> = 0.0 to 1.5, α<sub>chain,I</sub> = −0.7 to 0.7 |
| Fig. 3*F* | α<sub>recip,E</sub> = 0.0, α<sub>conv,E</sub> = 1.0, α<sub>div,E</sub> = 1.0, α<sub>chain,E</sub> = −0.5 to 0.5 |
|  | α<sub>recip,I</sub> = 0.0, α<sub>conv,I</sub> = 0.0 to 1.5, α<sub>div,I</sub> = 0.0, α<sub>chain,I</sub> = 0.0 |
| Fig. 3*G* | α<sub>recip,E</sub> = 0.0, α<sub>conv,E</sub> = 1.0, α<sub>div,E</sub> = 1.0, α<sub>chain,E</sub> = −0.5 to 0.5 |
|  | α<sub>recip,I</sub> = 0.0, α<sub>conv,I</sub> = 0.5, α<sub>div,I</sub> = 0.5, α<sub>chain,I</sub> = −0.25 to 0.25 |
| **Figure 4** |  |
| Fig. 4*A* (circle) | α<sub>recip,E</sub> = 0.0, α<sub>conv,E</sub> = 0.0, α<sub>div,E</sub> = 0.0, α<sub>chain,E</sub> = 0.0 |
|  | α<sub>recip,I</sub> = 0.02, α<sub>conv,I</sub> = 0.0, α<sub>div,I</sub> = 0.0, α<sub>chain,I</sub> = 0.0 |
| Fig. 4*A* (square) | α<sub>recip,E</sub> = 0.13, α<sub>conv,E</sub> = 1.20, α<sub>div,E</sub> = 1.13, α<sub>chain,E</sub> = 0.71 |
|  | α<sub>recip,I</sub> = 0.02, α<sub>conv,I</sub> = 0.0, α<sub>div,I</sub> = 0.0, α<sub>chain,I</sub> = 0.0 |
| Fig. 4*A* (diamond) | α<sub>recip,E</sub> = 0.13, α<sub>conv,E</sub> = 1.20, α<sub>div,E</sub> = 1.13, α<sub>chain,E</sub> = 0.71 |
|  | α<sub>recip,I</sub> = −0.09, α<sub>conv,I</sub> = 1.04, α<sub>div,I</sub> = 1.09, α<sub>chain,I</sub> = 0.60 |
| Fig. 4*A* (triangles) | Same connectivity parameters as Fig. 4A (square); 75% of the inhibitory neurons received an additional current of 0–120 pA. |

For the experiments shown in figure 2, we are only varying one of the *α* parameters while keeping the others fixed. The variable parameter is chosen at regular intervals over a particular range, as shown in table 3.

For the first three rasters of figure 3, we have chosen three particular cases for the example spiking data. The numbers in the table show the calculated *α* values from the networks. For the remaining four figures showing the scatter plots in figure 3, the parameters were chosen uniformly in a fixed range that is shown in the table. For figure 3 D and E, we had a total of 462 and 443 non-random excitatory and inhibitory matrices generated. For the figure 3 F and G, we had 100 matrices for each of the cases. Their combinations were taken for the simulation experiments.

**Figure 3:**
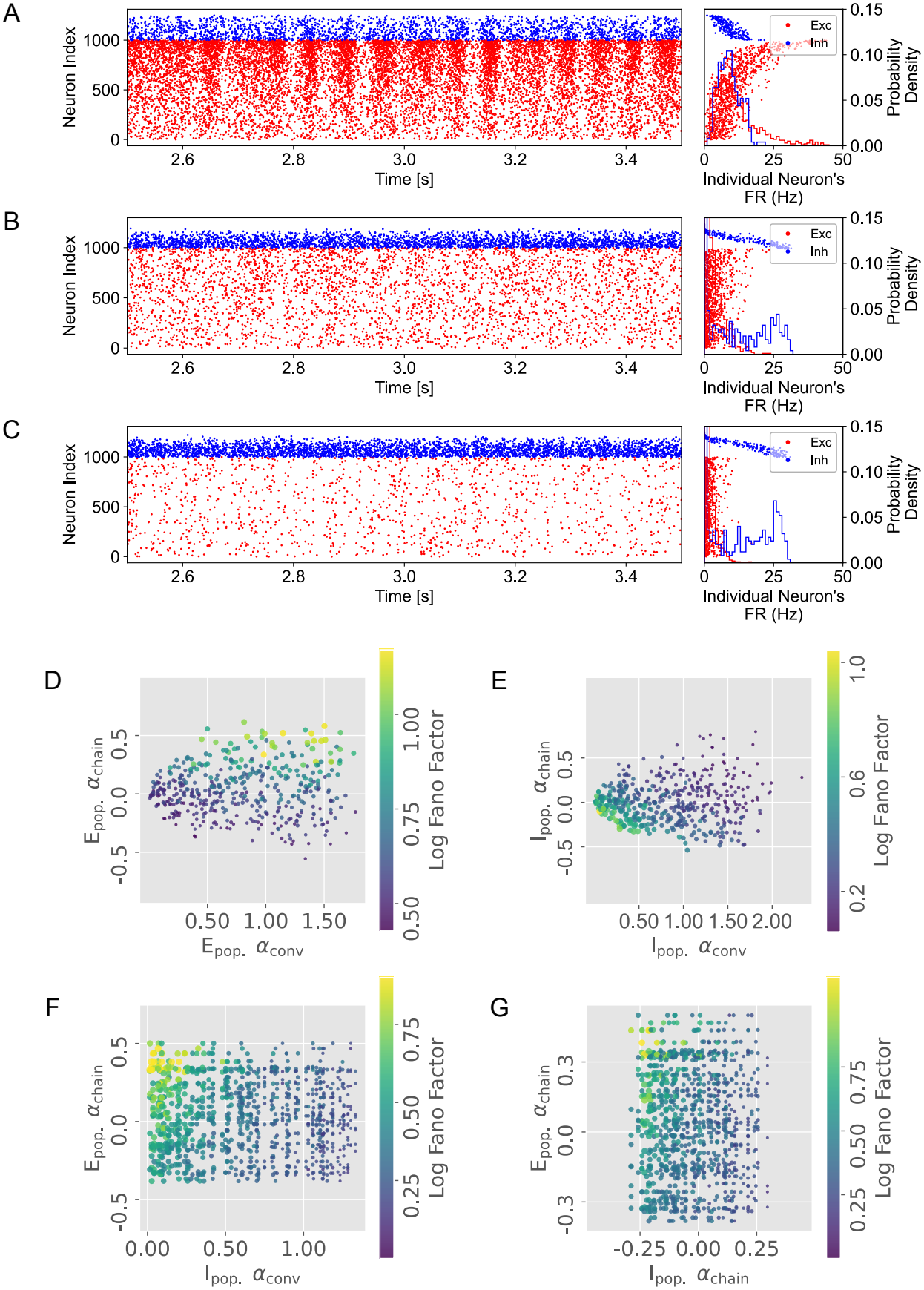
Interaction between second order motifs in excitatory (E) and inhibitory (I) populations. **(A–C)** Raster plots for different fractions of motifs in the E and I populations. Neurons were sorted in ascending order according to their in-degree. **A** E-population had chain motifs while the I-population had random connectivity. The right panel shows the distribution of firing rate of individual neurons using scatter alongside the histogram of the time-averaged firing rate distribution across neurons. **B** E-population had chain motifs while I-population had convergent motifs. **C** Both E- and I-populations had chain motifs. The quenching of synchrony was more prominent when chain motifs were present in the I-populations than when convergent motifs were present (compare panels B and C). (**D**) Scatter plot showing how the Fano factor (FF) of the E-population varies with the prevalence of chain and convergent motifs in the E-population network, while the I-population network was set to random. Each dot is one simulation, with color showing the FF, and the size is proportional to the firing rate (FR) of the E-population. The *α* values in the E-population were uniformly randomly selected from a specified range, whereas the *α* values in the I-population were all zero.**(E)**Same as in **D**, but here the scatter plot shows how FF (of E-population) varies with the prevalence of chain and convergent motifs in the I-population network, while the E-population network was set to random.**(F)**Scatter plot showing the effect of chain motifs in E-population and convergent motifs in I-population, while the prevalence of other motifs was set to a fixed value. (**G**) Same as in **F**, but the x-axis now refers to chain motifs in the I-population, and the y-axis remains chain motifs in the E-population, while the prevalence of all other motifs was fixed.

For Figure 4A, we used three representative network conditions: E-random/I-random, E-chain/I-random, and E-chain/I-chain. The calculated *α* values for these networks are listed in Table 3. The current-driven conditions used the same E-chain/I-random connectivity while an additional current (on top of the 1000 *Hz* Poisson input) was injected into a randomly selected 75% of the inhibitory neurons.

**Figure 4:**
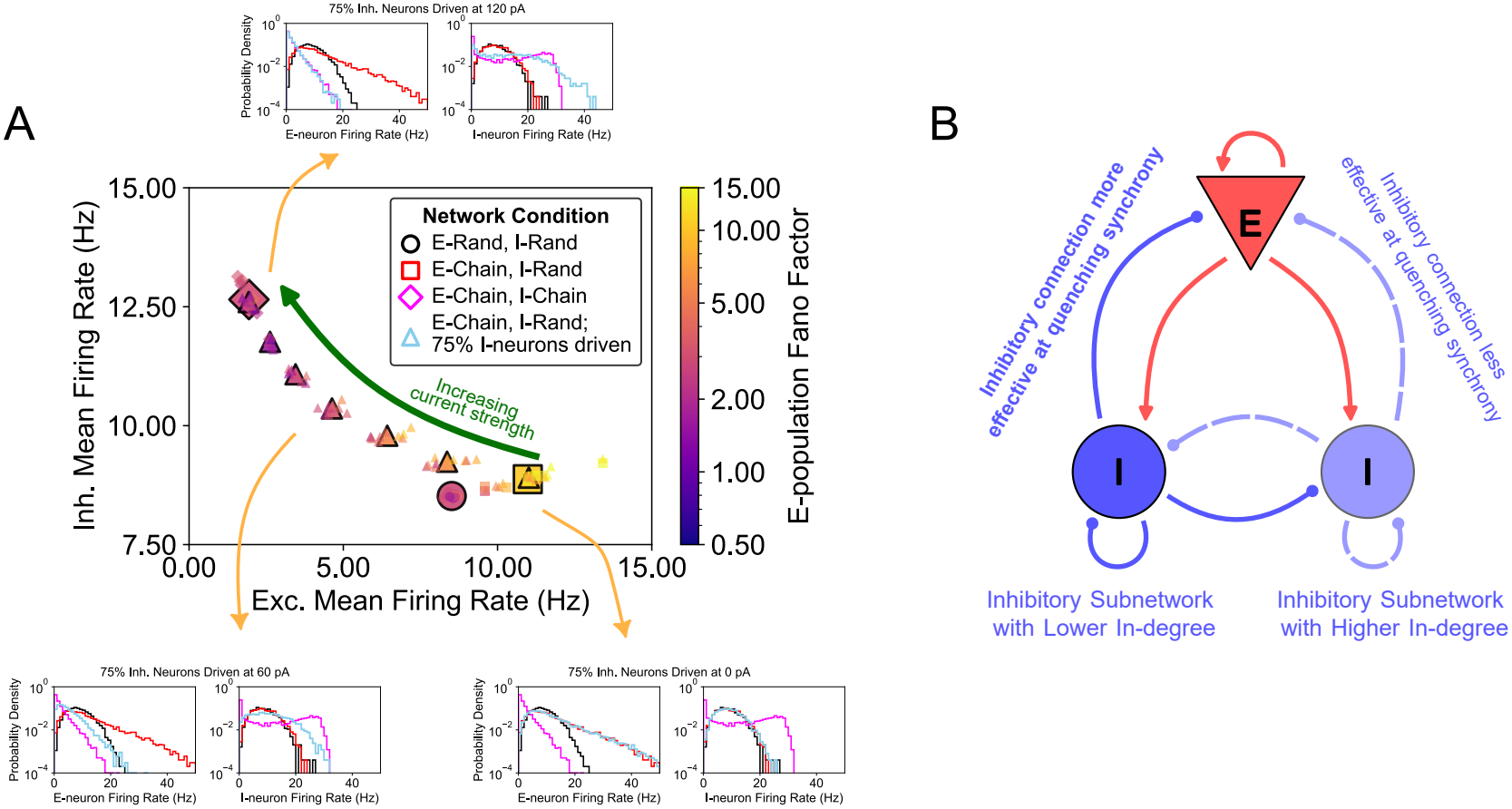
Increasing heterogeneity in the inhibitory firing-rate distribution suppresses network synchrony. (**A**) State-space representation of the EI-network dynamics. The axes show the mean firing rates of the excitatory (Exc.) and inhibitory (Inh.) populations, while marker color indicates the Fano factor of the excitatory population on a logarithmic scale. Circles, squares, and diamonds denote the E-random/I-random, E-chain/I-random, and E-chain/I-chain network conditions, respectively. Triangles denote E-chain/I-random networks in which randomly selected 75% of the inhibitory neurons received an additional current. Within each simulation condition, the same current amplitude was applied to every selected neuron; across conditions, the amplitude was varied from 0 to 120 pA in increments of 20 pA. Large markers with black outlines indicate means across 10 independent trials, whereas smaller markers indicate individual trials. The insets show the excitatory and inhibitory single-neuron firing-rate distributions (pooled across the 10 trials) for representative current amplitudes of 0 *pA*, 60 *pA*, and 120 *pA*. Black, red, and magenta curves denote the E-random/I-random, E-chain/I-random, and E-chain/I-chain conditions, respectively, while cyan curves denote the current-driven E-chain/I-random condition. (**B**) Schematic of the proposed mechanism by which chain motifs in the inhibitory population suppress synchrony. Inhibitory motifs broaden the inhibitory in-degree distribution, producing sub-populations with lower and higher recurrent inhibitory in-degree. This heterogeneity causes unequal recruitment of inhibitory neurons and distributes inhibition across neurons with different levels of recurrent inhibition, thereby reducing coherent excitatory population activity.

### Analysis of simulated network activity

#### Fano Factor

To measure the synchrony in the network, the Fano factor (FF) of the population spike count was calculated as follows:

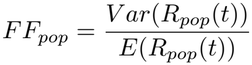

Here, *R*_*pop*_(*t*) is the averaged spike count across all neurons in either the E or I population for a given rectangular window around the time point *t*. For the results shown in our work, the window width in Fano factor calculations was chosen to be 5 *ms*. If the spiking activity is Poissonian in nature, then the Fano factor will be equal to 1. Whereas the presence of synchrony in the network leads to an increase in the numerator *V ar*(*R*_*pop*_(*t*)). Hence, Fano factors *≥* 1 are expected for synchronous activity.

### Pairwise Correlation Values

To further characterize the structure of neural activity, we computed the pairwise Pearson correlation coefficients between the binned spike count time series of all neurons. For any two neurons *i* and *j*, let *S*_*i*_(*t*) and *S*_*j*_(*t*) denote their respective spike count time series obtained by binning with a window of 100 ms and step size of 20 ms. The correlation coefficient between these two neurons is defined as:

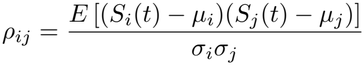

where *µ*_*i*_ and *σ*_*i*_ are the mean and standard deviation of the time series *S*_*i*_(*t*), given by:

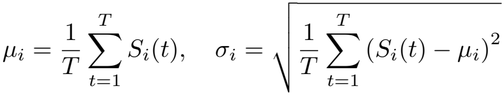

Here, *T* is the total number of time bins. This correlation coefficient *ρ*_*ij*_ quantifies the degree of linear dependence between the activity of neurons *i* and *j*.

### Participation Ratio

Another metric that was used to measure the amount of synchrony in the network is the participation ratio or the dimensionality of the covariance matrix. In order to calculate the participation ratio (*D*), a time series with the binned spike counts is calculated for each of the neurons in the network. For the binning, the window width was chosen to be 30 *ms* and step size was chosen as 10 *ms*. Suppose that we have the binned spike series for the *i*^*th*^ neuron to be *S*_*i*_(*t*). Then the covariance matrix is given by:

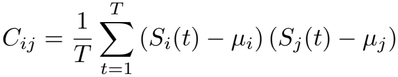

Here,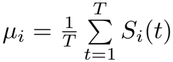is the mean spike count of neuron *i* over *T* time bins. If the matrix ***C*** as eigenvalues *λ*_1_, *λ*_2_, …, *λ*_*N*_, then the participation ratio is given by:

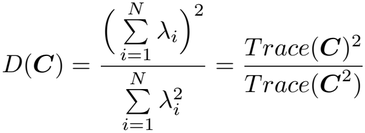

### Effective Rank

The connectivity matrices used here are sparse (all simulations have *p* = 0.1 unless stated otherwise). The probability of these sparse matrices being of full rank is very close to 1, even in the presence of row and column similarities. We used a measure we call effective rank, which is an entropy-based measure of the rank of the matrix. It measures how uniformly the singular values of the connectivity matrix ***W*** are distributed. Suppose *s*_*i*_(*≥* 0) are the singular values of ***W***, then we can define the normalized singular values as:

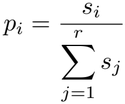

The entropy of the distribution is given by:

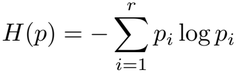

And, the effective rank of the matrix is defined as the exponent of the entropy:

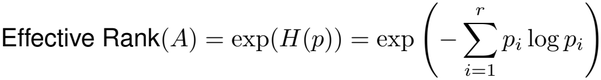

For a matrix of size *N × N*, the effective rank will be *N* only if all the singular values are equal. As the singular values distribution gets more skewed, the effective rank also becomes lower.

### Simulation and data analysis tools

All simulations were performed in BRIAN [25]. The simulations and analyses were performed using custom python scripts. Simulation code is available on GitHub (https://github.com/ArcherFerrata/Biswas-and-Kumar-2025).

## Data availability

Simulation code is available on GitHub (https://github.com/ArcherFerrata/Biswas-and-Kumar-2025). There is no experimental data used in the manuscript. Simulation data can be easily recreated by running the code.

## Results

To understand how non-random motif counts shape the dynamics (firing rate and synchrony), we systematically varied the reciprocal, convergent, divergent, and chain motifs (Figure 1A) in a network of excitatory and inhibitory neurons (Figure 1C). We refer to these motifs as di-synaptic or second-order motifs because to construct networks with such motifs, we need to consider two connections simultaneously (see Methods). We considered a network in which leaky-integrate-fire neurons were connected via static synapses. We chose this setting with a simple neuron and synapse model so as to extract the effects of the network structure. **Later, we also show that the key synchrony-promoting and synchrony-quenching effects persist in networks composed of Exponential Integrate-and-Fire (EIF) neurons, indicating that these results are not specific to the precise spike-generation mechanism of the LIF model. Unless explicitly stated otherwise, all simulations and analyses presented below were performed using networks composed of LIF neurons**.

### Second order connectivity motifs alter degree distributions and correlations

Global network properties such as effective rank, degree distribution, and correlation between in- and out-degree are closely related to network activity dynamics (firing rate and synchrony) [12, 22, 23, 13, 7, 26, 17, 15, 20]. Therefore, before delving into the dynamics of networks with second order motifs we first investigated how the introduction of such motifs affected the global network properties.

We used the SONETs algorithm [22] to vary the number of second order connectivity motifs in the network (see Methods). SONETs allowed us to introduce di-synaptic motifs (second-order motifs) of 2/3 neurons without affecting the average connection probability. Maintaining a constant average connection probability is essential, as it isolates the effects of motif structure from those of network sparsity. For instance, the reciprocal motif tends to make the network connectivity matrix more symmetric (by definition of the motif, Figure 2Ai). Systematic increase in the number of reciprocal motifs did not affect the rank of the connectivity matrix (Figure 2Bi) and variance of the in- and out-degree distribution (Figure 2Ci, green traces), it increased the degree correlation (Figure 2Ci, orange trace).

Increasing the convergent or divergent motifs resulted in horizontal or vertical stripes in the connectivity matrix, respectively (Figure 2Aii-iii). That is, with more convergent motifs, some neurons received more input as compared to the ER network. Similarly, with divergent motifs, some neurons sent out more output projections as compared to the ER network. This increased the variance of in-degree (for convergent motifs) and out-degree (for divergent motifs) (Figure 2Cii-iii, green traces, square and circular markers respectively). In contrast to the reciprocal motif, convergent and divergent motifs reduced the rank of the connectivity matrix (Figure 2Bii-iii) without affecting the degree correlations (Figure 2Cii-iii, orange trace).

Introduction of chain motifs results in the formation of plaid patterns, with both vertical and horizontal stripes in the connectivity matrix (Figure 2Aiv). To increase chain motifs, we fixed the amount of divergent and convergent motifs. Therefore, increasing the amount of chain motifs did not affect the variance in the degree distributions (Figure 2Civ, green traces), but it increased the correlation between in- and out-degree (Figure 2Civ, orange trace). The effective rank is not affected by the increase in chain motifs either (Figure 2Biv).

Thus, changing the reciprocal, convergent, divergent, and chain motifs affects the higher-order properties of the connectivity matrix, such as degree variance, degree correlations, and effective rank.

### Only convergent or chain motifs among excitatory neurons increase synchrony

Next we characterized how second order connectivity motifs affected the activity of EI networks. First, we focused on the excitatory population and systematically varied the prevalence of reciprocal, convergent, divergent, and chain motifs while keeping the inhibitory population connectivity random. The overall connection probability was maintained at *p* = 0.1.

We found that only convergent and chain motifs strongly promoted network synchrony (e.g. Figure 3A). Networks enriched with these two motifs showed an increase in synchrony (Fano factor) and firing rate (Figure 3D). By contrast, reciprocal and divergent motifs had little to no effect on either metric (Figure S2,S4). Increasing chain motifs in particular shifted the dynamics toward more correlated activity. These results are consistent with previous results which have argued that shared connectivity causes synchrony in EI networks [27]. Motifs in the excitatory population thus form a structural mechanism to obtain high synchrony while keeping the average connectivity low. To obtain a similar level of high synchrony in an Erdos-Renyi network (without motifs), we would require much higher average connectivity (Figure S1,S2).

### Effect of convergent and chain motifs among inhibitory neurons

The emergence of synchrony due to di-synaptic motifs among excitatory neurons is at odds with the experimental data, which showed that cortical activity is only weakly correlated [24]. Therefore, next, we asked how the synchrony induced by excitatory di-synaptic motifs could be quenched. Recent experimental data suggest that there may be di-synaptic motifs within the Inh. to Inh. connectivity [13]. Therefore, we hypothesize that di-synaptic motifs in the inhibitory population can quench the synchrony.

To test this hypothesis, first, we asked how motifs within the inhibitory population influence network activity, keeping excitatory connectivity random. As in the previous section, we systematically varied the prevalence of reciprocal, convergent, divergent, and chain motifs while keeping the excitatory population connectivity random. The overall connection probability was maintained at *p* = 0.1.

The results reveal a striking contrast to the case of excitatory population motifs. While convergent and chain motifs in the E-population enhanced synchrony and elevated firing rates, the same motifs in the I-population had the opposite effect (compare Figure 3A and B). An increase in inhibitory convergent and/or chain motifs reduced both the synchrony and the average excitatory firing rate (Figure 3E). By contrast, inhibitory reciprocal and divergent motifs had little effect on the network activity (Figure S3,S5). Thus, these results suggest that inhibitory motifs could quench the synchrony induced by convergent/chain motifs in the excitatory population.

### Inhibitory motifs can counteract synchrony generated by excitatory motifs

Next, to test our hypothesis, we introduced di-synaptic motifs within excitatory and within inhibitory populations. Inter-population connections were kept random. In such a network the opposing effects of excitatory and inhibitory motifs became more evident. When motifs were present in both populations, excitatory motifs (convergent and chain) increased network firing rates and synchrony, while the inhibitory motifs (convergent and chain) quenched these effects (Figure 3C, F, G).

To illustrate this, we compared networks in which excitatory chain motifs were systematically increased while inhibitory motifs were varied in parallel. In both cases, inhibitory convergent and chain motifs reduced the Fano factor, offsetting the synchrony that would otherwise emerge due to motifs in the excitatory population. Notably, chain motifs in the inhibitory population were more effective suppressors of synchrony than convergent motifs (Figure 3G).

These results highlight that inhibitory motifs play an important role in desynchronizing the network activity: by counteracting excitatory motifs, they can effectively bring down the overall synchrony level of the EI network. Overall, the presence of motifs in both the excitatory and inhibitory populations expands the dynamical range of network activity in terms of both firing rate and synchrony.

### Desynchronization persists at fixed firing rate

In the previous sections, we observed that firing rate and synchrony were often correlated, raising the question of whether inhibitory motifs quench synchrony only because they reduce excitatory population activity. To disentangle these effects, we compared networks with different motif structures while adjusting the input firing rates (same for E and I populations) in order to maintain the average firing rate at a fixed value.

In both EI network without motifs and with only E-motifs, firing rate and synchrony were highly correlated. In general, for the same average output rate, networks with E-motifs showed higher synchrony than networks without any di-synaptic motifs (compare red/blue and black dots in Figure S7A). Moreover, chain motifs resulted in higher synchrony than convergent motifs (compare red and blue dots in Figure S7A).

By contrast, in networks with both E- and I-motifs, firing rate was no longer correlated with synchrony, and for the same firing rates these networks showed almost no synchrony (Figure S7A, cyan, green, magenta dots). Inhibitory chain motifs were particularly more effective at reducing the Fano factor, outperforming those with inhibitory convergent motifs (compare magenta and cyan/green dots in Figure S7A). Thus, we can argue that the synchrony-reducing effect of inhibitory motifs is not just a byproduct of a decrease in firing rates.

### Heterogeneous inhibitory activity mediates the synchrony-quenching effect of inhibitory motifs

Inclusion of motifs altered the higher-order connectivity of the network (Figure 2). In particular, convergent and chain motifs increased the variance of the inhibitory in-degree distribution, causing inhibitory neurons to receive different amounts of recurrent inhibition. We therefore hypothesized that the synchrony-quenching effects of these motifs arise from heterogeneous firing of the inhibitory neurons.

To test whether heterogeneity in firing rate of inhibitory neurons was sufficient to quench synchrony, we considered a network with only E-motifs (chain) and I neurons were connected randomly. Such networks show synchrony (3A). In this network we randomly selected 75% inhibitory neurons and injected extra input so as to increase their firing rate and thereby create heterogeneity in the firing rates of inhibitory neurons. We varied the amplitude of the additional current from 0 pA to 120 pA in increments of 20 pA. We then compared the resulting network states in terms of the mean excitatory firing rate, mean inhibitory firing rate, and synchrony (Fano factor) within the excitatory population (Figure 4A).

Without additional current, the network with E-chain/I-random connectivity exhibited high excitatory activity and strong excitatory population synchrony. As the current delivered to the selected inhibitory neurons increased, the network progressively shifted toward lower excitatory activity, higher inhibitory activity, and reduced excitatory population synchrony (Figure 4A). At the largest current amplitude ( 120 pA), the network state was similar to that observed when chain motifs were introduced into the inhibitory connectivity.

For each additional input configuration we also estimated the distribution of single-neuron firing-rate (Figure 4A, insets). As the amplitude of the additional current increased, the inhibitory firing-rate distribution broadened and developed a bimodal structure (for the maximal input we used). This redistribution of inhibitory activity was accompanied by a shift of the excitatory firing-rate distribution toward lower values and a reduction in excitatory population synchrony. Consequently, the firing-rate distributions increasingly approached that generated by introducing chain motifs into the inhibitory network, even though the inhibitory connectivity itself remained random. These experiments demonstrate that a heterogeneous recruitment of otherwise randomly connected inhibitory neurons is sufficient to reproduce the synchrony-quenching effect associated with inhibitory motifs.

Thus the synchrony quenching effect of I-motifs can be understood in the following manner (Figure 4B): Motifs in the inhibitory population broaden the inhibitory in-degree distribution, thereby generating subpopulations that receive different levels of recurrent inhibition. This results in a wide distribution of inhibitory firing rate. In particular, inhibitory neurons that received fewer recurrent inhibitory connections maintain high firing rate. Such neurons are crucial for quenching synchrony as they are more effective in canceling the network fluctuations created by excitatory-motifs [28].

### The Synchrony-Quenching Effects Persist in EIF Networks

We next asked whether these effects of di-synaptic motifs depended on the neuronal model. To test this, we repeated the simulations using exponential integrate-and-fire (EIF) neurons. We observed the same qualitative effects as in networks with LIF neurons. Adding chain motifs to the excitatory population increased its firing rate and promoted synchronized population activity. Adding convergent or chain motifs to the inhibitory population suppressed this synchrony. Inhibitory chain motifs produced the stronger suppression (Figure S6).

Thus, these effects are not restricted to LIF neurons. Emergence of synchrony due to E-motifs (chain and convergent) and quenching of synchrony due to I-motifs is a property of the network which arises due the distribution of in- and out-degree of neurons.

## Discussion

Here, we investigate how second order motifs affect the network structure and dynamics, in particular synchronization. It is by now well established that excitatory neurons have an overabundance of second-order motifs as compared to an Erdő s–Rényi type random network [8, 29, 9]. Consistent with previous work [22, 15], we show that convergent and chain motifs in the excitatory population tend to synchronize network activity. The high synchrony, characteristics of network with E-motifs, is not often observed in *in vivo* in healthy states; therefore, it is important to identify possible means by which network activity can be desynchronized in the presence of E-motifs.

Our key result here is that second order motifs in inhibitory neurons reduce both synchrony and firing rate. We show that I-motifs can counter the synchrony produced by E-motifs and make the network activity physiologically more realistic. Our findings also suggest that, in the context of the model and dynamical regime explored, reciprocal and divergent motifs do not play a significant role in shaping the network dynamics.

### From second order motifs to large scale structure

At the network structure level one of the key insights from our work is that the second order motifs manifest themselves at the large network scales in the form of variance of degree distribution, correlations between in- and out-degree, and overall rank of the connectivity matrix (Figure 2). Each second order motif leaves its unique signature, e.g. convergent motifs increase in-degree variance while divergent motifs increase out-degree variance. Reciprocal and chain motifs tend to increase in- and out-degree correlations and so on. Unlike the low-rank of the connectivity matrix, it is these large-scale changes in the degree variance and covariances that shape the network dynamics. As our results show, that low rank in the connectivity matrix is not sufficient to guarantee low-dimensional activity (e.g. as argued by Mastrogiuseppe et al. [17]). In our model, second-order motifs make the network low rank, but excitatory and inhibitory interactions can and do cancel the low-rank effects and make the activity asynchronous or high-dimensional.

### Role of in-degree distribution in controlling synchrony

If there is high in-degree variance, then some neurons receive more recurrent input than others. The neurons that receive relatively fewer recurrent inputs are only weakly affected by the network activity, and their activity is largely governed by external inputs. If the external input to the low in-degree neurons is asynchronous (as is the case in our model), these neurons will tend to decorrelate the activity and vice versa. Similarly, when there is high variance in the out-degree, neurons with high out-degree control the dynamics. Roxin et al. [20] reported similar results to ours using a network made up of only I-neurons having high variance in in-degree or out-degree distributions. We complement that study by showing that a biologically plausible way to achieve high variance in the in/out-degree is through second order motifs.

The decorrelation of network activity is not simply due to the fact that low in-degree inhibitory neurons (which have a high firing rate) reduced the excitatory firing rate. In our study, we systematically varied the prevalence of motifs in both the E and I populations, allowing us to observe how firing rate and synchrony change smoothly across different motif counts. Our results show that, even after controlling for differences in firing rate, networks containing I-motifs remain substantially less synchronous than networks without I-motifs (Figure S7A)

### Model predictions

Given that the network activity is largely asynchronous [30, 31] and there are second order motifs among excitatory neurons [8, 9, 32, 13, 10], there should be similar di-synaptic motifs among inhibitory neurons. There is already partial support for this prediction [13].

We also predict that only convergent and chain type motifs among excitatory neurons affect synchrony in a network, while other motifs, such as reciprocal and divergent motifs, do not play a significant role in shaping the network dynamics. The count of the I-motifs shall be high enough to counterbalance the synchrony induced by excitatory motifs. At the core of the desynchronization mechanism is the wide in-degree distribution of inhibitory neurons. In the absence of inhibitory motifs, we would predict that in cortical networks, inhibitory neurons should have a wide inhibitory in-degree distribution. In the cortical microcircuit, SST+ neurons are not recurrently connected unlike the PV positive neurons [33], and they could also control the synchrony. In general, our model predicts that stimulation (activation/inactivation) of inhibitory neurons with low recurrence will have a much greater impact on network response than other inhibitory neurons.

Our simulations show that different motif configurations produce distinct single-neuron firing-rate distributions. Firing-rate heterogeneity may therefore provide an activity-based signature of the degree heterogeneity generated by second-order motifs, although it cannot uniquely identify the underlying motif structure.

### Functional significance

What we have shown is that while E-motifs induce synchrony, I-neurons can quench that (even completely). However, quenching depends on the activity of inhibitory neurons, which receive relatively fewer recurrent inhibitory connections. Thus, these di-synaptic motifs give the network a large dynamical range in the space of firing rate and synchrony with sparse average connectivity. To achieve the same range of firing rate and synchrony in Erdő s–Rényi type networks, one would require far higher connection probability (Figure S1).

While it is likely that the di-synaptic motifs are a simple consequence of neuron morphologies [10], motif statistics can also be shaped by learning and synaptic plasticity. Irrespective, the connectivity heterogeneity created by motifs forms an interesting way to control brain dynamics, especially correlations.

### Beyond motifs

In computational models, it is not trivial to study the effect of a certain circuit motif as we need to carefully control its prevalence without affecting other properties. This can only be done for motifs of 2-3 neurons. Beyond that, there is a combinatorial explosion, and it is not feasible to construct networks with specific motif statistics. However, as our results show, di-synaptic motifs manifest as variance and covariance of in- and out-degree. Networks with specific degree variance and covariance are relatively easy to construct, and their effect on network dynamics is easier to study. With the advent of viruses that cross a synapse [34] and complete reconstruction of network connectivity [35], it is now possible to measure the in- and out-degree statistics.

### Limitations

To get insights into the role of di-synaptic motifs on network activity, we had to make several assumptions. Here, we discuss three key assumptions that may have implications for the validity of our predictions. First, we primarily used a simple leaky integrate-and-fire (LIF) model and tested the robustness of our principal findings in selected simulations using exponential integrate-and-fire (EIF) neurons. Although the EIF results indicate that the synchrony-promoting and synchrony-quenching effects are not specific to the spike-generation mechanism of the LIF model, both models remain simplified point-neuron descriptions. Neuronal properties not represented here, including adaptation, bursting, and other cell-type-specific nonlinearities, may interact with motif-induced connectivity heterogeneity and generate dynamics beyond those observed in our networks. Second, we have only considered a network with one excitatory and one inhibitory population. Networks in different brain regions have multiple types of excitatory and inhibitory neurons, which follow type specific connectivity rules. The effect of the introduction of motifs in such a network might be drastically different from what we have observed here. But, the systematic study of the effects of motifs also becomes much more difficult as there are many more combinations in which the motifs can be introduced in the different populations in such a network. But, it becomes difficult to characterize such connectivity rules that include multiple neuron types, even when considering motifs made up of only three neurons, as shown in [23, 13]. However, our insight that in- and out-degree statistics are more crucial than exact motif structure, provides a strategy to study heterogeneous networks. Third, we have assumed that all synaptic weights of a given type have the same strength. Synaptic weight distribution might dilute the effect of motifs because, in such networks, the two connections in a di-synaptic motif will have unequal strengths. However, this needs to be determined to what extent the effect of motifs can be overridden by weight distributions. Short-term dynamics of synapses could also modulate the effect of di-synaptic motifs on the network dynamics. These assumptions raise interesting new questions that should be addressed in future studies.

Despite simplifying assumptions, we show that non-random features can expand the range of the network activity and provide additional means to control brain dynamics.

## Acknowledgments

We thank Rajat Mishra for helpful discussions and feedback.

## Author Contribution

AK and AB conceived and designed the study. AB performed all the simulations and data analysis. AB wrote the first draft of the manuscript. AK edited the manuscript and supervised project.

## Funding declaration

This project was funded in parts by Swedish Research Council (StratNEURO SRA and SeRC SRA) and Institute of Advanced Studies University of Strasbourg, France, Fellowship to AK.

## Supplementary Figures

**Figure S1:**
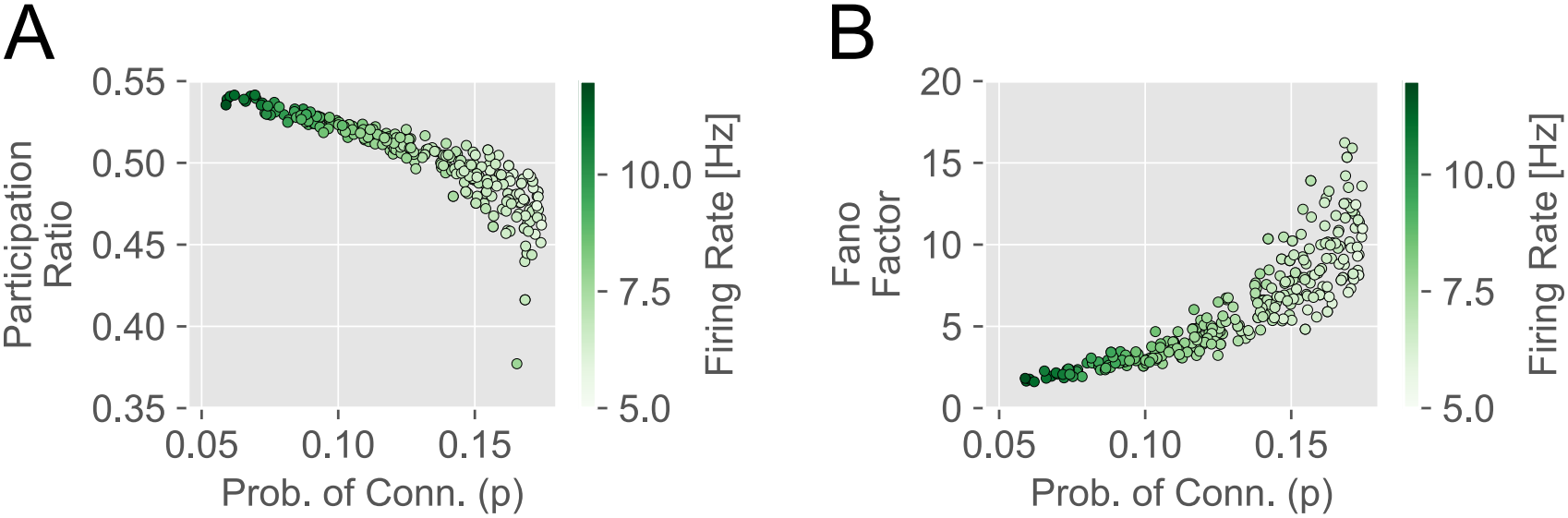
Effects of connection probability on participation ratio and Fano factor in random EI networks. (**A**) Shows the calculated participation ratio (dimensionality) from the E-population activity for different EI networks with random connectivity (ER network) in both the population and having different probabilities of connection (*p*). A total of 300 ER (Erdő s–Rényi) connectivity matrices were generated with parameter *p* sampled uniformly between 0.05 and 0.175. The networks were stimulated with the Poisson input as described in the methods section. Each scatter point denotes one simulation run with the x-position as the calculated probability of connection from the whole network and the y-position as the calculated participation ratio from the E-population activity. (**B**) Same as that of the plot in **A**, except that the y-position of the scatter points is now the Fano factor calculated from the E-population activity. In both plots, the color represents the firing rate calculated from the E-population.

**Figure S2:**
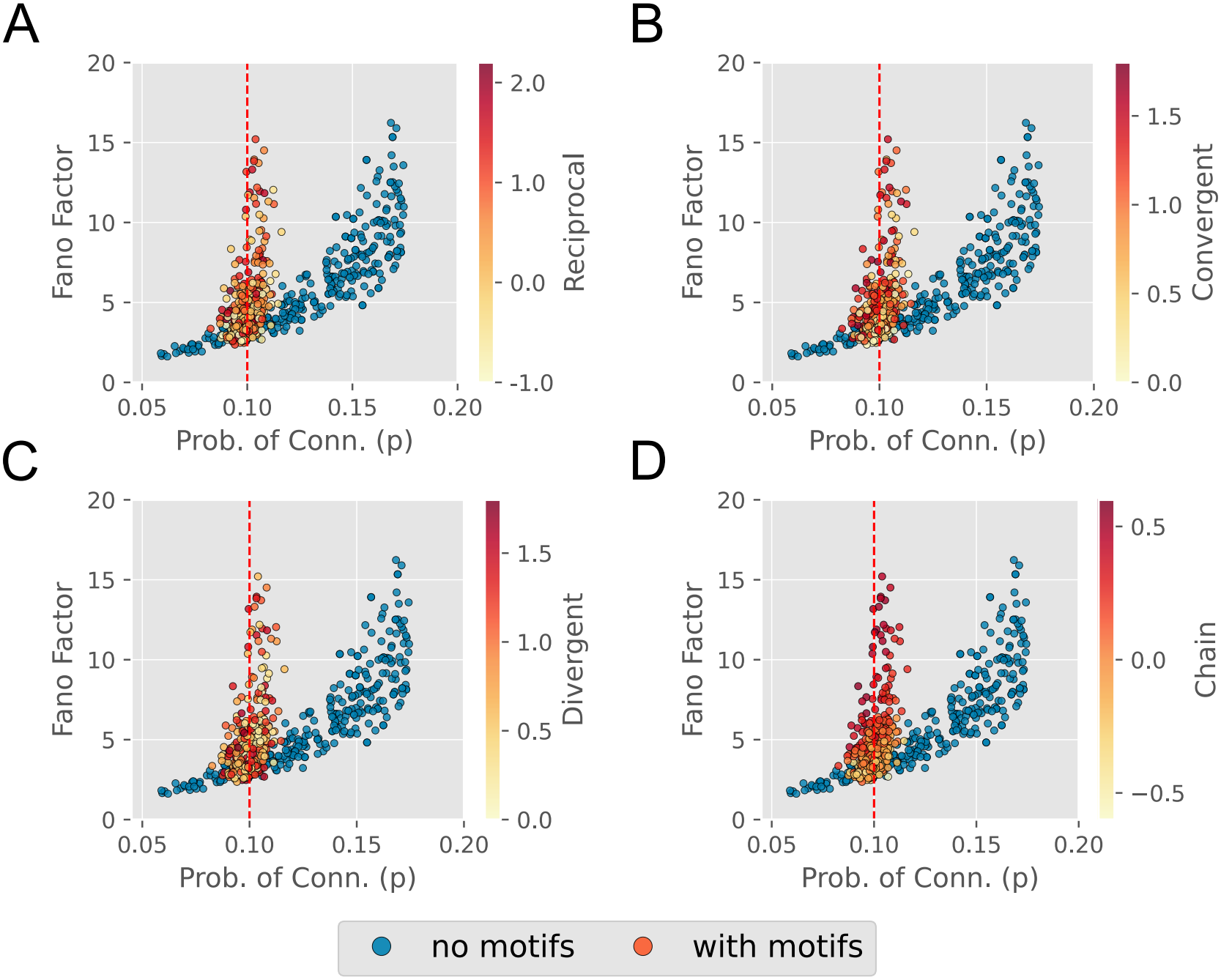
Effects of excitatory population motifs and connection probability on the Fano factor. The four scatter plots have in common the blue dots that show how the Fano factor changes with the probability of connection (*p*) in an ER EI network, same as in figure S1. For the other scatter dots with the colorbar, they belong to the simulations that were plotted in 3(D) and later in S4. Motifs were introduced into the E-population network with *α* values randomly chosen in a given range, and the I-population network was kept random. The Fano factor (of E-population) v/s measured sparsity *p* (of E-population) is plotted. In each of the four plots, only the coloring of the dots corresponding to the simulation runs with motifs in the E-population is different. The colors in the four plots show the calculated *α*_*recip*_(in **A**), *α*_*conv*_(in **B**), *α*_*div*_(in **C**), and *α*_*chain*_(in **D**) from the E-population connectivity matrix.

**Figure S3:**
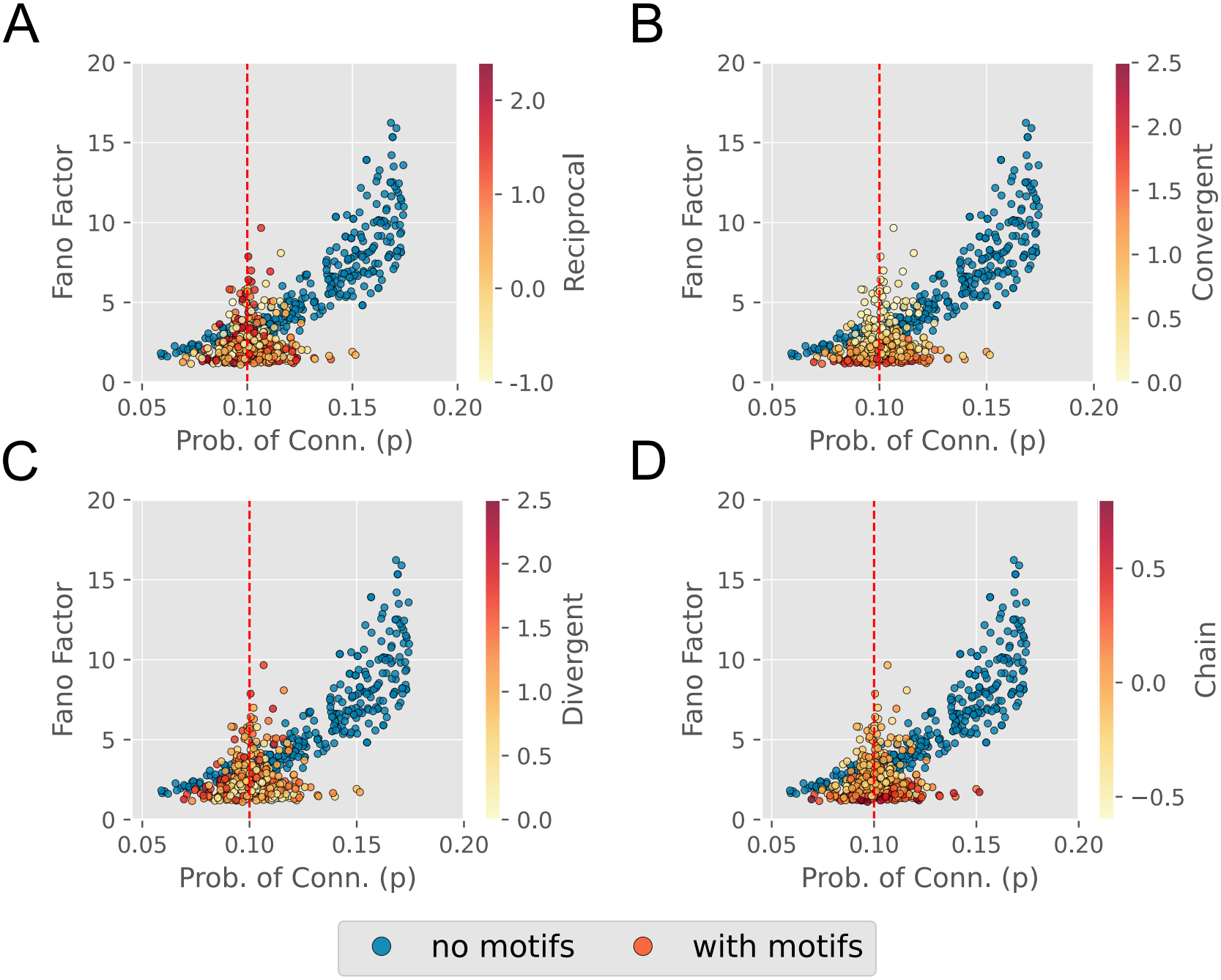
Effects of inhibitory population motifs and connection probability on the Fano factor. The four scatter plots have in common the blue dots that show how the Fano factor changes with the probability of connection (*p*) in an ER EI network, same as in figure S1. For the other scatter dots with the colorbar, they belong to the simulations that were plotted in 3(E) and later in S5. Motifs were introduced into the I-population network with *α* values randomly chosen in a given range, and the E-population network was kept random. The Fano factor (of E-population) v/s measured sparsity *p* (of I-population) is plotted. In each of the four plots, only the coloring of the dots corresponding to the simulation runs with motifs in the I-population is different. The colors in the four plots show the calculated *α*_*recip*_(in **A**), *α*_*conv*_(in **B**), *α*_*div*_(in **C**), and *α*_*chain*_(in **D**) from the I-population connectivity matrix.

**Figure S4:**
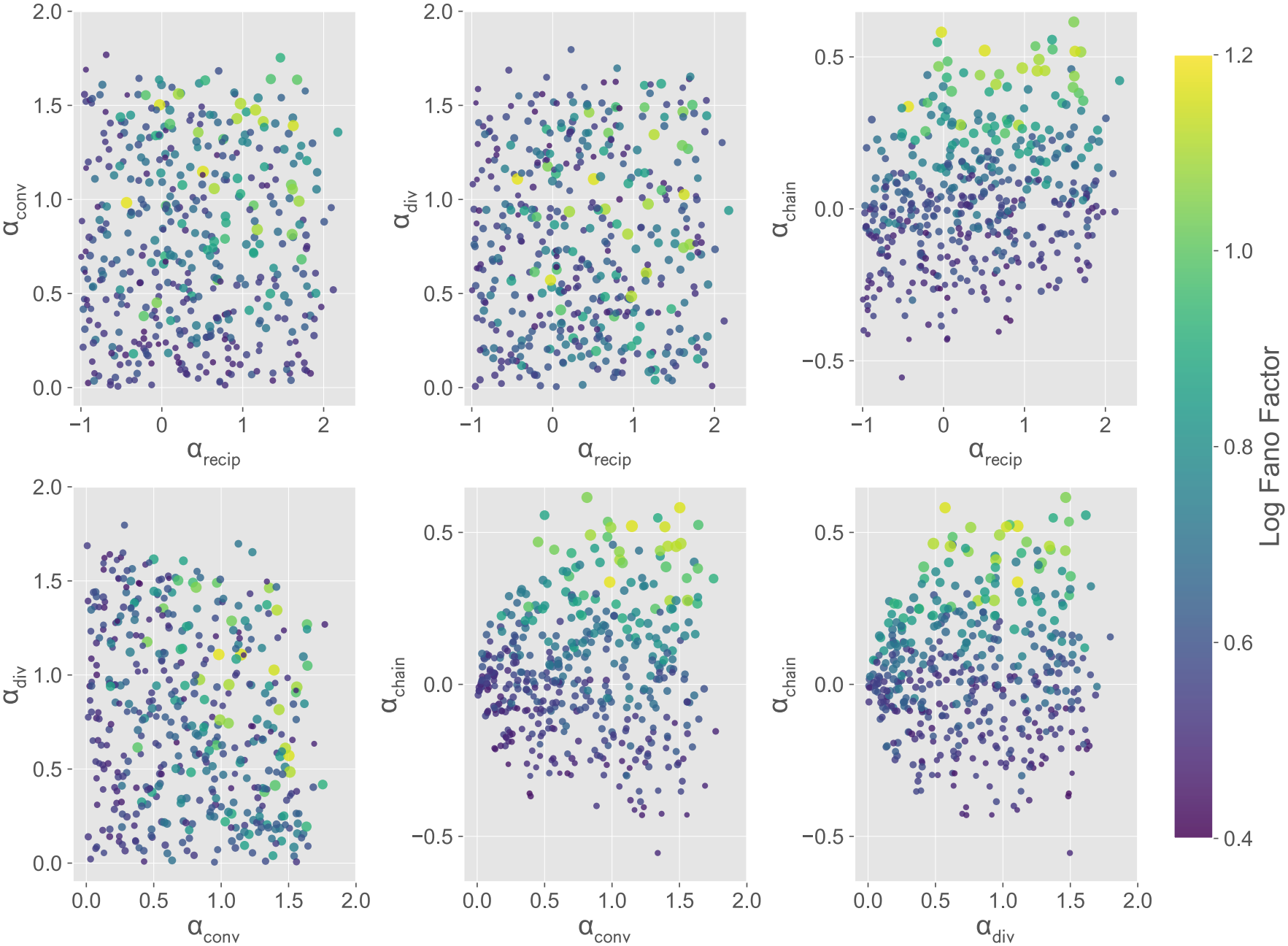
Effects of excitatory population motif parameters on Fano factor and firing rate. Each of the six subplots belongs to the same set of simulations. In this set of simulations, the *α* values in the E-population were randomly selected from a specified range, whereas the *α* values in the I-population were all set to zero. For all simulations, the *p* = 0.1. Six subplots are shown, corresponding to pairwise combinations of the four *α* parameters: *α*_*recip*_, *α*_*conv*_, *α*_*div*_, and *α*_*chain*_ calculated from the E-population network. The color shows the Fano factor calculated from the activity of the E-population. The size of the dots is set proportional to the firing rate of the E-population. The figure 3(D) shows one particular subplot among them: the chain motifs (*α*_*chain*_) vs convergent (*α*_*conv*_) motifs of the E-population.

**Figure S5:**
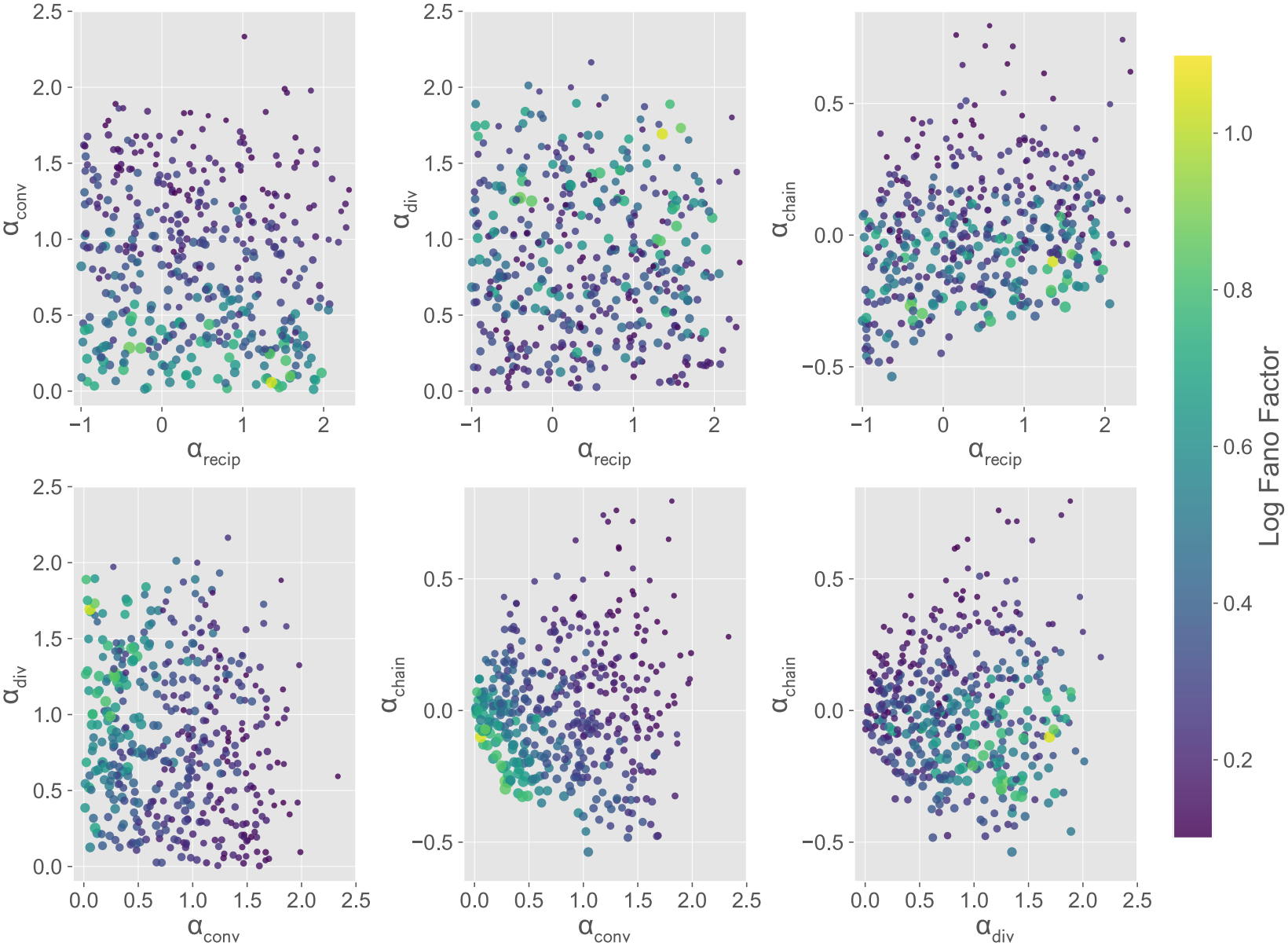
Effects of inhibitory population motif parameters on Fano factor and firing rate. Similar to figure S4, but in this set of simulations, the *α* values in the I-population were randomly selected from a specified range, whereas the *α* values in the E-population were all set to zero. For all simulations, the *p* = 0.1. Six subplots are shown, corresponding to pairwise combinations of the four *α* parameters: *α*_*recip*_, *α*_*conv*_, *α*_*div*_, and *α*_*chain*_ calculated from the I-population network. The color shows the Fano factor calculated from the activity of the E-population. The size of the dots is set proportional to the firing rate of the E-population. The figure 3(E) shows one particular subplot among them: the chain motifs (*α*_*chain*_) vs convergent (*α*_*conv*_) motifs of the I-population.

**Figure S6:**
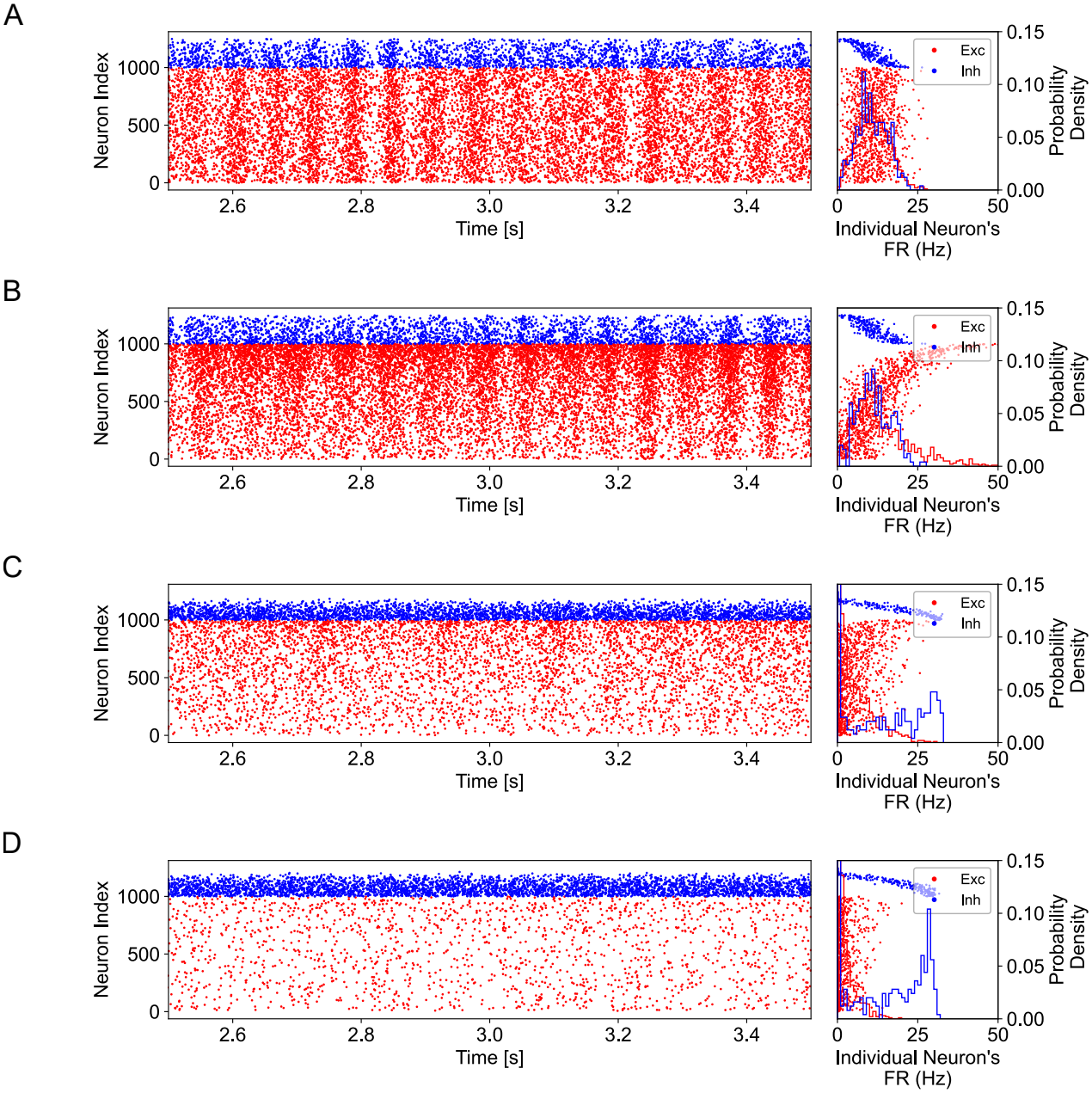
The effects of excitatory and inhibitory second-order motifs persist in networks of exponential integrate-and-fire (EIF) neurons. (**A–D**) Raster plots and single-neuron firing-rate distributions for four network conditions. Neurons were sorted in ascending order according to their in-degree. The panels to the right of each raster show the firing rates of individual neurons together with the distribution of time-averaged firing rates across neurons. (**A**) Both the E- and I-populations had random connectivity.(**B**) The E-population contained chain motifs, while the I-population had random connectivity. (**C**) The E-population contained chain motifs, while the I-population contained convergent motifs. (**D**) Both the E- and I-populations contained chain motifs. As in the leaky integrate-and-fire network, introducing motifs into the inhibitory population suppressed the synchrony generated by excitatory chain motifs, with stronger suppression observed for inhibitory chain motifs than for inhibitory convergent motifs (compare panels C and D).

**Figure S7:**
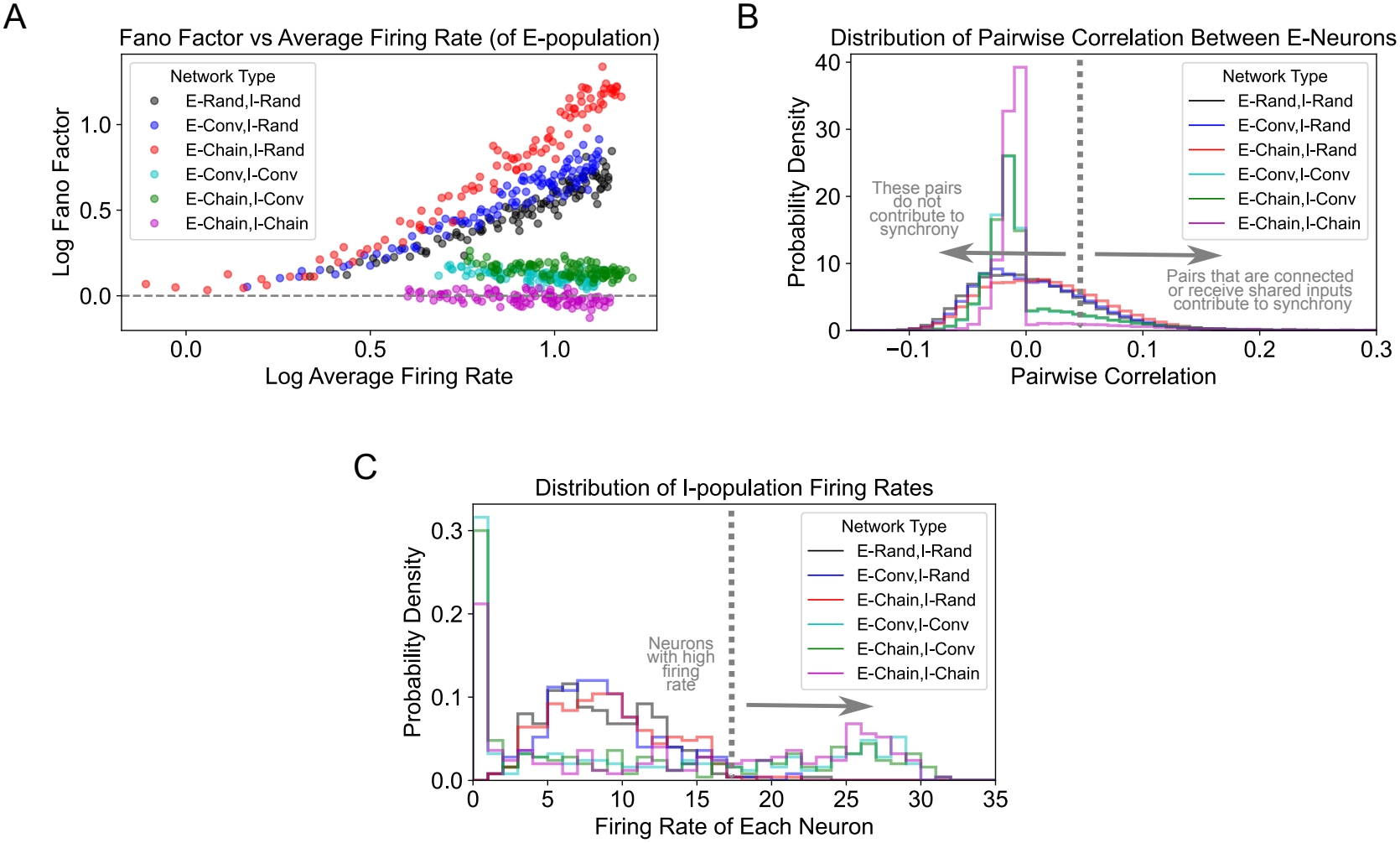
Distributions of correlations and firing rates, and synchrony–rate relation for different motif configurations in E and I populations. (**A**) Scatter plot showing the relation between the synchrony (E-population’s Fano Factor) and the E-population’s average firing rate when driven with Poisson inputs of different rates. (**B**) Distribution of pairwise correlation values across all the pairs of neurons in the E-population for different types of motifs in the E and I populations, which is shown in the legends. (**C**) Distribution of firing rate across individual inhibitory neurons. The different colors, which are kept the same across the three panels, are for different types of motifs in the E and I population, as shown in the legends.

## Notes

### Competing Interest Statement

The authors have declared no competing interest.

### Summary of Updates

Revised the Figure 4 to elaborate. the role of in-degree distribution of inhibitory neurons in controlling the synchrony.

## References

[1] O. Sporns and J. D. Zwi. “The Small World of the Cerebral Cortex”. In: Neuroinformatics 2.2 (2004), pp. 145–162. DOI: 10.1385/ni:2:2:145.

[2] S. Pajevic and D. Plenz. “Efficient Network Reconstruction from Dynamical Cascades Identifies Small-World Topology of Neuronal Avalanches”. In: PLoS Computational Biology 5.1 (Jan. 2009). Ed. by O. Sporns, e1000271. DOI: 10.1371/journal.pcbi.1000271.

[3] J. M. Beggs and D. Plenz. “Neuronal Avalanches in Neocortical Circuits”. In: The Journal of Neuro-science 23.35 (Dec. 2003), pp. 11167–11177. DOI: 10.1523/jneurosci.23-35-11167.2003.

[4] J. M. Beggs and D. Plenz. “Neuronal Avalanches Are Diverse and Precise Activity Patterns That Are Stable for Many Hours in Cortical Slice Cultures”. In: The Journal of Neuroscience 24.22 (June 2004), pp. 5216–5229. DOI: 10.1523/jneurosci.0540-04.2004.

[5] X. Xu, N. D. Olivas, T. Ikrar, T. Peng, T. C. Holmes, Q. Nie, et al. “Primary visual cortex shows laminar-specific and balanced circuit organization of excitatory and inhibitory synaptic connectivity”. In: The Journal of physiology 594.7 (2016), pp. 1891–1910.

[6] P. Schnepel, A. Kumar, M. Zohar, A. Aertsen, and C. Boucsein. “Physiology and impact of horizontal connections in rat neocortex”. In: Cerebral Cortex 25.10 (2015), pp. 3818–3835.

[7] E. Gal, R. Perin, H. Markram, M. London, and I. Segev. “Neuron Geometry Underlies Universal Network Features in Cortical Microcircuits”. In: (May 2019). DOI: 10.1101/656058.

[8] S. Song, P.J. Sjöström, M. Reigl, S. Nelson, and D. B. Chklovskii. “Highly Nonrandom Features of Synaptic Connectivity in Local Cortical Circuits”. In: PLoS Biology 3.3 (Mar. 2005). Ed. by K. J. Friston, e68. DOI: 10.1371/journal.pbio.0030068.

[9] R. Perin, T. K. Berger, and H. Markram. “A synaptic organizing principle for cortical neuronal groups”. In: Proceedings of the National Academy of Sciences 108.13 (Mar. 2011), pp. 5419–5424. DOI: 10.1073/pnas.1016051108.

[10] D. Udvary, P. Harth, J. H. Macke, H.-C. Hege, C. P. de Kock, B. Sakmann, et al. “The impact of neuron morphology on cortical network architecture”. In: Cell Reports 39.2 (Apr. 2022), p. 110677. DOI: 10.1016/j.celrep.2022.110677.

[11] A. Lin, R. Yang, S. Dorkenwald, A. Matsliah, A. R. Sterling, P. Schlegel, et al. “Network statistics of the whole-brain connectome of Drosophila”. In: Nature 634.8032 (Oct. 2024), pp. 153–165. DOI: 10.1038/s41586-024-07968-y.

[12] V. Pernice, B. Staude, S. Cardanobile, and S. Rotter. “How structure determines correlations in neuronal networks”. In: PLoS computational biology 7.5 (2011), e1002059.

[13] D. Dahmen, S. Recanatesi, X. Jia, G. K. Ocker, L. Campagnola, S. Seeman, et al. “Strong and localized recurrence controls dimensionality of neural activity across brain areas”. In: (Nov. 2020). DOI: 10.1101/2020.11.02.365072.

[14] L. Tiberi, D. Dahmen, and M. Helias. Hidden connectivity structures control collective network dynamics. 2023. DOI: 10.48550/ARXIV.2303.02476.

[15] Y. Shao, D. Dahmen, S. Recanatesi, E. Shea-Brown, and S. Ostojic. “Impact of Local Connectivity Patterns on Excitatory-Inhibitory Network Dynamics”. In: PRX Life 3.2 (May 2025). DOI: 10.1103/prxlife.3.023008.

[16] S. Recanatesi, G. K. Ocker, M. A. Buice, and E. Shea-Brown. “Dimensionality in recurrent spiking networks: Global trends in activity and local origins in connectivity”. In: PLOS Computational Biology 15.7 (July 2019). Ed. by J. Diedrichsen, e1006446. DOI: 10.1371/journal.pcbi.1006446.

[17] F. Mastrogiuseppe and S. Ostojic. “Linking Connectivity, Dynamics, and Computations in Low-Rank Recurrent Neural Networks”. In: Neuron 99.3 (Aug. 2018), 609–623.e29. DOI: 10.1016/j.neuron.2018.07.003.

[18] L. Cimeša, L. Ciric, and S. Ostojic. “Geometry of population activity in spiking networks with low-rank structure”. In: PLOS Computational Biology 19.8 (Aug. 2023). Ed. by P. E. Latham, e1011315. DOI: 10.1371/journal.pcbi.1011315.

[19] M. Beiran, A. Dubreuil, A. Valente, F. Mastrogiuseppe, and S. Ostojic. “Shaping Dynamics With Multiple Populations in Low-Rank Recurrent Networks”. In: Neural Computation 33.6 (May 2021), pp. 1572–1615. DOI: 10.1162/neco_a_01381.

[20] A. Roxin. “The Role of Degree Distribution in Shaping the Dynamics in Networks of Sparsely Connected Spiking Neurons”. In: Frontiers in Computational Neuroscience 5 (2011). DOI: 10.3389/fncom.2011.00008.

[21] D. Q. Nykamp, D. Friedman, S. Shaker, M. Shinn, M. Vella, A. Compte, et al. “Mean-field equations for neuronal networks with arbitrary degree distributions”. In: Physical Review E 95.4 (Apr. 2017). DOI: 10.1103/physreve.95.042323.

[22] L. Zhao, B. Beverlin, T. Netoff, and D. Q. Nykamp. “Synchronization from Second Order Network Connectivity Statistics”. In: Frontiers in Computational Neuroscience 5 (2011). DOI: 10.3389/fncom.2011.00028.

[23] Y. Hu, J. Trousdale, K. Josić, and E. Shea-Brown. “Motif statistics and spike correlations in neuronal networks”. In: Journal of Statistical Mechanics: Theory and Experiment 2013.03 (Mar. 2013), P03012. DOI: 10.1088/1742-5468/2013/03/p03012.

[24] A. S. Ecker, P. Berens, G. A. Keliris, M. Bethge, N. K. Logothetis, and A. S. Tolias. “Decorrelated Neuronal Firing in Cortical Microcircuits”. In: Science 327.5965 (Jan. 2010), pp. 584–587. DOI: 10.1126/science.1179867.

[25] M. Stimberg, R. Brette, and D. F. Goodman. “Brian 2, an intuitive and efficient neural simulator”. In: eLife 8 (Aug. 2019). Ed. by F. K. Skinner, e47314. DOI: 10.7554/eLife.47314.

[26] H. Setareh, M. Deger, C. C. H. Petersen, and W. Gerstner. “Cortical Dynamics in Presence of Assemblies of Densely Connected Weight-Hub Neurons”. In: Frontiers in Computational Neuroscience 11 (June 2017). DOI: 10.3389/fncom.2017.00052.

[27] E. Shea-Brown, K. Josić, J. de la Rocha, and B. Doiron. “Correlation and Synchrony Transfer in Integrate-and-Fire Neurons: Basic Properties and Consequences for Coding”. In: Physical Review Letters 100.10 (Mar. 2008). DOI: 10.1103/physrevlett.100.108102.

[28] T. Tetzlaff, M. Helias, G. T. Einevoll, and M. Diesmann. “Decorrelation of neural-network activity by inhibitory feedback”. In: (2012).

[29] E. Gal, M. London, A. Globerson, S. Ramaswamy, M. W. Reimann, E. Muller, et al. “Rich cell-type-specific network topology in neocortical microcircuitry”. In: Nature Neuroscience 20.7 (June 2017), pp. 1004–1013. DOI: 10.1038/nn.4576.

[30] W. Softky and C. Koch. “The highly irregular firing of cortical cells is inconsistent with temporal integration of random EPSPs”. In: The Journal of Neuroscience 13.1 (Jan. 1993), pp. 334–350. DOI: 10.1523/jneurosci.13-01-00334.1993.

[31] A. Renart, J. de la Rocha, P. Bartho, L. Hollender, N. Parga, A. Reyes, et al. “The Asynchronous State in Cortical Circuits”. In: Science 327.5965 (Jan. 2010), pp. 587–590. DOI: 10.1126/science.1179850.

[32] S. Rieubland, A. Roth, and M. Häusser. “Structured Connectivity in Cerebellar Inhibitory Networks”. In: Neuron 81.4 (Feb. 2014), pp. 913–929. DOI: 10.1016/j.neuron.2013.12.029.

[33] C. K. Pfeffer, M. Xue, M. He, Z. J. Huang, and M. Scanziani. “Inhibition of inhibition in visual cortex: the logic of connections between molecularly distinct interneurons”. In: Nature neuroscience 16.8 (2013), pp. 1068–1076.

[34] Q. Liu, Y. Wu, H. Wang, F. Jia, and F. Xu. “Viral tools for neural circuit tracing”. In: Neuroscience bulletin 38.12 (2022), pp. 1508–1518.

[35] “Functional connectomics spanning multiple areas of mouse visual cortex”. In: Nature 640.8058 (2025), pp. 435–447.

